# *DNAJC13* influences responses of the extended reward system to conditioned stimuli: a genome-wide association study

**DOI:** 10.1101/2022.05.03.490395

**Authors:** Jens Treutlein, Karolin E. Einenkel, Bernd Krämer, Swapnil Awasthi, Stephan Ripke, Oliver Gruber

**Author notes:** correspondence should be send to Professor Dr. Oliver Gruber, Address: Voßstraße 4, 69115 Heidelberg, Germany. shared first authorship (Jens Treutlein, Karolin E. Einenkel).

## Abstract

Reward system dysfunction is implicated in the pathogenesis of major psychiatric disorders. We conducted a genome-wide association study (GWAS) to identify genes that influence activation strength of brain regions within the extended reward system in humans. A large homogeneous sample of 214 participants was genotyped and underwent functional magnetic resonance imaging (fMRI). All subjects performed the ‘desire-reason dilemma’ (DRD) paradigm allowing systematic investigation of systems-level mechanisms of reward processing in humans. As a main finding, we identified the single nucleotide variant rs113408797 in the DnaJ Heat Shock Protein Family Member C13 gene (*DNAJC13*, alias RME-8), that strongly influenced the activation of the ventral tegmental area (VTA; p = 2.50E-07) and the nucleus accumbens (NAcc; p = 5.31E-05) in response to conditioned reward stimuli. Moreover, haplotype analysis assessing the information across the entire *DNAJC13* locus demonstrated an impact of a five-marker haplotype on VTA activation (p = 3.21E-07), which further corroborates a link between this gene and reward processing. The present findings provide first direct empirical evidence that genetic variation of *DNAJC13* influences neural responses within the extended reward system to conditioned stimuli. Further studies are required to investigate the role of this gene in the pathogenesis and pathophysiology of neuropsychiatric disorders.

## INTRODUCTION

The genetic architecture of major psychiatric disorders is known to be highly polygenic with a substantial amount being contributed by common genetic variation. This polygenic picture, where thousands of common variants of small effect size account for a proportion of the genome-wide contribution to a phenotype, is the case for most complex traits. By means of genome-wide association studies (GWAS), that query the whole genome for such genetic variation, polygenic features can be investigated. However, the biological mechanisms underlying the functional contribution of these variants are by far less well understood.

The endophenotype concept in psychiatry attempts to fill the gap between genetic contributions to a disorder and the elusive disease processes [1]. This approach is used to subdivide behavioral symptoms into more stable phenotypes for measuring phenotypic variation. It provides opportunities to identify the traits underlying clinical phenotypes and could facilitate the identification of susceptibility genes of mental disorders [1,2]. Variations in magnetic resonance imaging (MRI) signals can be considered as endophenotypes as it was shown that even healthy first-degree relatives of diseased persons show similar trait expressions as the affected ones [3,4]. Therefore, imaging genetics can shed light on genes, which increase the susceptibility and vulnerability to develop a psychiatric disorder. Among the neurofunctional systems that contribute to the etiology of psychiatric disorders, the extended dopaminergic reward system is most relevant and can be assessed by circuit-specific functional neuroimaging paradigms.

The reward system in the brain is based on a neural circuitry including regions of the mesolimbic dopamine (DA) system, in particular the ventral tegmental area (VTA) and the nucleus accumbens (NAcc) [5]. In a functional MRI (fMRI) study, we found that bipolar patients showed a reduced bottom-up responsiveness of the ventral striatum (vStr) and a disturbed top-down control of the mesolimbic reward system by prefrontal brain regions while performing a specific reward paradigm, the ‘desire-reason dilemma’ (DRD) paradigm [6]. The DRD paradigm assesses the neural mechanisms underlying reward processing and active reward dismissal in favor of a long-term goal [7]. Additionally, recent fMRI studies showed that patients with schizophrenia exhibited abnormal subcortical reward processing [8-10] and associated alterations in functional connectivity within the salience network and reward regions [11].

In the present study, we aimed to identify genetic variation that influences the reactivity of the mesolimbic extended reward system in response to conditioned reward stimuli by GWAS. To this end, a homogenous group of healthy adults underwent fMRI while performing the DRD paradigm. We expected that individuals that carry relevant genetic variants, particularly at previously implicated dopaminergic gene loci, show changes in subcortical reward processing.

## MATERIALS AND METHODS

### Subjects

Participants were recruited by advertisements in intern online student networks and local newsletters. Exclusion criteria were past or present psychiatric disorders according to ICD-10, a positive family history of psychiatric disorders, substance abuse during the last month, cannabis abuse during the last two weeks, mental retardation, dementia, neurological or metabolic diseases, and pregnancy in women. All participants were Caucasians with European ancestry.

Imaging and performance data as well as quality-controlled GWAS genotypes were available for 214 participants (mean age ±SD: 24.12 years ±2.46 years, 128 females). Exclusion criteria were failure of fMRI data acquisition, fMRI motion artifacts and/or less than 70% correct answers across all task conditions as well as missing genetic information and ancestry outliers, respectively. All participants had a higher school qualification.

Every subject provided written informed consent after the study procedure had been fully explained. The study was carried out in accordance with the latest version of the Declaration of Helsinki and was approved by the local ethics committee.

### Genotyping

DNA of all participants was isolated from saliva, which was collected using Oragene·DNA devices (OG-500) with standardized protocols from DNA Genotek (DNA Genotek, Ottawa, Ontario, Canada). Genome-wide single nucleotide polymorphism (SNP) genotyping was performed using Illumina OmniExpress Genotyping BeadChips according to the manufacturer’s protocols (Illumina, Inc., San Diego, California, USA). Prior to association analysis, ancestry outliers were identified by principal component analysis and excluded (Supplementary Methods, Supplementary Figure 1). Details on the imputation procedure are provided in Supplementary Methods. Single marker and haplotype association analyses for 214 individuals were performed on autosomal SNPs. Of 7872128 variants contained in the imputed dataset, 133395 variants were removed due to missing genotype data, 205824 variants due to Hardy-Weinberg exact test, and 3034535 variants due to allele frequency threshold, leaving 4498374 variants for genetic association analysis after main filters. Quantil-quantil (QQ) plots and inflation factor lambda were calculated as described in the Supplementary Methods and are shown in Supplementary Figure 2.

### Experimental Paradigm

Prior to the fMRI measurement, participants underwent a training session outside the scanner. Initially, the training started with an operant conditioning task. Squares with eight different colors, which were repeated 20 times each, were presented on a monitor in a randomized sequence. Subjects were instructed to respond to each of the stimuli by button press (left button: accept, right button: reject) with their right index and middle finger, respectively. Button choice was free and subjects were encouraged to explore the stimulus-response-reward contingencies. By doing so, subjects were conditioned to associate two colors (red and green) with an immediate reward (i.e., bonus of 10 points), four other colors with a neutral outcome (0 points) and two colors with a punishment (i.e., loss of 10 points). The goal of this operant conditioning task was to establish stimulus-response-reward contingencies for the next phase of the experiment.

During the second phase of the training, subjects were familiarized with a modified version of the DRD paradigm, a sequential forced-choice task described by Diekhof, et al. ^12^ and firstly introduced by Diekhof and Gruber ^7^. Subjects had to perform blocks of four or eight trials. At the beginning of every block, two targets (two different neutral colors) were shown. In the following, four or eight colored squares were presented one after another. Two different types of blocks had to be performed. In the first type, the ‘Desire Context’ (DC), subjects were allowed to accept the two previously conditioned reward stimuli in addition to the two target colors in order to win bonus points (i.e., +10 points). In the second type, the ‘Reason Context’ (RC), only the two target colors had to be accepted, but the conditioned reward stimuli had to be rejected in order to successfully pursue the goal to receive 50 points at the end of the block. Failure to choose the right button or to press the correct button timely led to an immediate termination of the current task block, zero outcome and the feedback ‘goal failure’. The task context always changed after two consecutive blocks and was indicated by a cue. For detailed information concerning task timing see Diekhof, et al. ^12^.

At the day after the training session, the participants performed 40 task blocks of the DRD paradigm over the course of two fMRI runs. All points gained during the experiment in the MR scanner were cashed into real money.

### fMRI Data Acquisition

Imaging was performed on a 3T MRI scanner (Magnetom TRIO, Siemens Healthcare, Erlangen, Germany) equipped with a standard eight-channel phased-array head coil. First, a T1-weighted anatomical data set with 1 mm isotropic resolution was obtained. Parallel to the anterior commissure– posterior commissure line, 31 axial slices were acquired in ascending acquisition order (slice thickness = 3 mm; interslice gap = 0.6 mm) using a T2*-sensitive echo-planar imaging (EPI) sequence (echo time 33 ms, flip angle 70°; field-of-view 192 mm, TR 1900 ms). In two functional runs, 185 volumes each were acquired. Subjects responded via button presses on a fiber optic computer response device (Current Designs, Philadelphia, Pennsylvania, USA), and stimuli were viewed through goggles (Resonance Technology, Northridge, California, USA). Triggering of the visual stimulation by the scanner impulse during functional data acquisition and generation of the stimuli was conducted through the Presentation® Software (Neurobehavioral Systems, Albany, California, USA).

### fMRI Data Preprocessing

Functional images were preprocessed with SPM12 (Statistical Parametric Mapping; https://www.fil.ion.ucl.ac.uk/spm/software/spm12/; SPM RRID:SCR_007037). The study design was event-related and only correctly answered trials were included in the analyses. Preprocessing involved realignment and unwarping, correction for slice-time acquisition differences by using the first slice as reference, normalization into standard stereotactic space (Montreal Neurological Institute [MNI]) and spatial smoothing with a 6 mm full-width at half-maximum (FWHM) isotropic Gaussian kernel filter.

### fMRI Statistics (first-level)

A general linear model (GLM) with 11 regressors (target, non-target and reward stimuli for the DC and RC condition, respectively, two context cues, the block cues and finally the block feedback for either successful goal completion or goal failure) was used for statistical analyses with SPM12 (Statistical Parametric Mapping; https://www.fil.ion.ucl.ac.uk/spm/software/spm12/). A vector representing the temporal onsets of stimulus presentation was convolved with canonical hemodynamic response function (hrf) to produce the predicted hemodynamic response to each experimental condition. Linear t-contrasts were defined by contrasting activation effects elicited by the conditioned stimuli in the DC relative to the implicit baseline to examine pure reward-related activation patterns.

### fMRI Phenotype Extraction

For the subsequent genetic association analyses, neuroimaging phenotypes were determined. The individual mean beta estimates from the *a priori* regions of interest (ROIs), namely the VTA and the NAcc, were extracted with the MarsBaR ROI toolbox for SPM [13]. Beta extraction for each participant was performed using predefined MNI coordinates of both brain regions emerging from previous publications: ±9 −21 −15 for the bilateral VTA, and ±12 12 −3 for the bilateral NAcc [12], respectively. To account for interindividual functional neuroanatomical variation, a box with a 3 × 3 × 3 mm^3^ dimension around the *a priori* MNI coordinates was applied to determine the individual maximum of reward-related brain activation within each subject.

### Genetic Association Analyses of Single Markers

Single marker association analyses were conducted via PLINK v2.00a2.3 (64-bit (24 Jan 2020); https://www.cog-genomics.org/plink/2.0/, PLINK RRID:SCR_001757), using linear model testing for quantitative traits (‘linear’ option) with imputed data. Further options were individual and genotyping call rates > 0.02, minor allele frequency > 0.05 and Hardy-Weinberg equilibrium (HWE) p > 0.05. The output of these analyses was used to retrieve the asymptotic p-values for t-statistic, effect-sizes (beta) and standard errors (SE). SNPs were annotated to genes as indicated by dbSNP [14](dbSNP database RRID:SCR_002338). Allele frequencies were calculated using the --freq command in PLINK v2.00a2.3.

We adapted our procedure to retrieve the most reliably associated chromosomal loci with our DA-related trait. Initially, sample bisection was used to narrow down the top ten associated SNPs for each phenotype and consequently yielding the most robustly associated and most plausible variants for the phenotype. For this purpose, we divided the full sample of 214 individuals into two half-samples consisting of 107 subjects each. Subsequently, a proportion of individuals, namely 22 subjects, was repeatedly exchanged between the two half-samples. After each replacement, a single-marker association analysis was conducted for both half-samples and for the best ten SNPs resulting from the initial whole-sample (214 individuals) single marker association analysis. SNPs that fulfilled the criterion p < 0.05 across all exchange steps in all half-samples were considered the most robustly associated SNPs. Then, the genes, to which these robustly associated SNPs were annotated, were subjected to a comprehensive literature search for an involvement in DA-related processes. This procedure resulted in three most plausible genes: *DNAJC13* (DnaJ Heat Shock Protein Family Member C13, synonym: Receptor-mediated endocytosis 8 [RME-8]), *IMMP2L* (Inner Mitochondrial Membrane Peptidase Subunit 2) and *SV2C* (Synaptic Vesicle Glycoprotein 2C).

### Genetic Association Analyses of Haplotypes

In order to further corroborate the most promising GWAS findings within the genes *DNAJC13, IMMP2L* and *SV2C*, we investigated if any of them received further support from haplotypic analysis spanning the entire gene. This strategy, to support a gene by haplotype analysis across the entire locus, was previously pursued by Stacey, et al. ^15^ and Jia, et al. ^16^. Among the three loci, only *DNAJC13* showed additional associative signals of gene-spanning haplotypes.

For haplotypic analysis of *DNAJC13*, the following procedure was used to minimize the marker number and to investigate the association: first, markers across the whole gene that fulfilled the criteria genotype call rate > 0.02, minor allele frequency > 0.2, and Hardy Weinberg equilibrium p > 0.05 were retrieved from the genotyped dataset. Boundaries for the *DNAJC13* gene were RefSeq +-20,000 bases on either side of the gene (chr3:132116346-132277876, hg19) (https://genome.ucsc.edu/; UCSC Genome Browser RRID:SCR_005780). This region comprises both *DNAJC13* RefSeq isoforms, NM_001329126 (isoform 1) and NM_015268 (isoform 2). The procedure resulted in ten markers (Supplementary Figure 4). Second, to further reduce the number of markers, block-by-block tag-markers were selected from the ten markers. For this purpose, haplotype blocks were generated with HaploView v.2.4 [17] (https://www.broadinstitute.org/haploview/haploview; Haploview RRID:SCR_003076) using the default block definition [18]. This procedure clustered nine of the ten markers into three blocks (markers 1-3 formed haplotype block 1, markers 5-6 haplotype block 2 and markers 7-10 haplotype block 3; marker 4 had no haplotype block affiliation; Supplementary Figure 4). We retained seven of the nine markers as they were displayed to constitute block-by-block tag-markers by HaploView. Third, we removed the entire intermediate block consisting of two block-by-block tag-markers, because haplotype association analyses of each of the three blocks separately indicated that five SNPs were sufficient to capture the association with the extent of activation of the right VTA (Supplementary Figure 4, Supplementary Table 3). Forth, association analysis of the five-marker haplotypes was conducted using PLINKv1.07 [19]. The linear regression option for quantitative outcomes (‘hap-linear’) was used to generate the asymptotic p-values, effect sizes (beta values) and frequencies of the haplotypes.

### Additional Whole-Brain Neuroimaging Analyses for *DNAJC13* Single Marker rs113408797

Whole-brain neuroimaging analyses of *DNAJC13* genetic subgroups were conducted to explore additional effects of this gene on brain regions outside VTA and NAcc. Using second-level analyses in SPM12 (Statistical Parametric Mapping; https://www.fil.ion.ucl.ac.uk/spm/software/spm12/) based on first-level single subjects contrast images, effects of rs113408797 were evaluated. For this purpose, a full factorial design with the levels ‘homozygous major allele carrier’ (n = 175), ‘homozygous minor allele carrier’ (n = 2) and ‘heterozygous allele carrier’ (n = 34) was applied to assess whole-brain genotypic effects on reward-related brain activation with a particular focus on the contrast ‘all minor allele carriers (homozygotes and heterozygotes) *versus* homozygous major allele carriers’. These analyses were conducted with a search criterion of p < 0.001, uncorrected, with subsequent FWE-correction for multiple comparisons as indicated in Table 2.

## RESULTS

### Genetic Association Analyses of Single Markers

Single marker analyses for the extent of activation generated several findings. Table 1 lists the detailed results for the top ten SNPs with the lowest p-values for each brain region. Regional association plots of the top ten SNPs are shown in Supplementary Figure 3. Association results up to a p-value of p < 1E-05 are displayed in Supplementary Table 1.

**Table 1.**
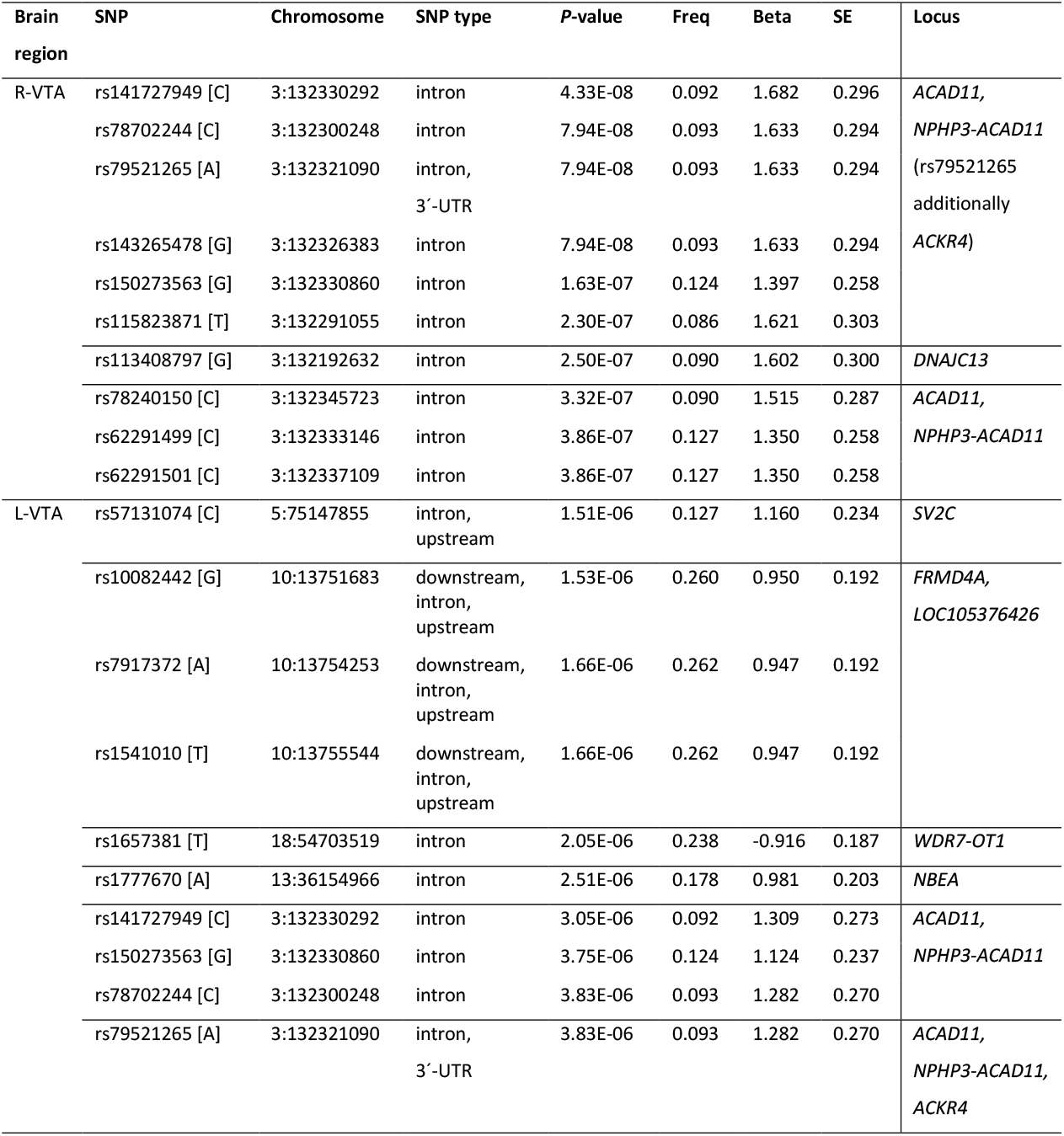
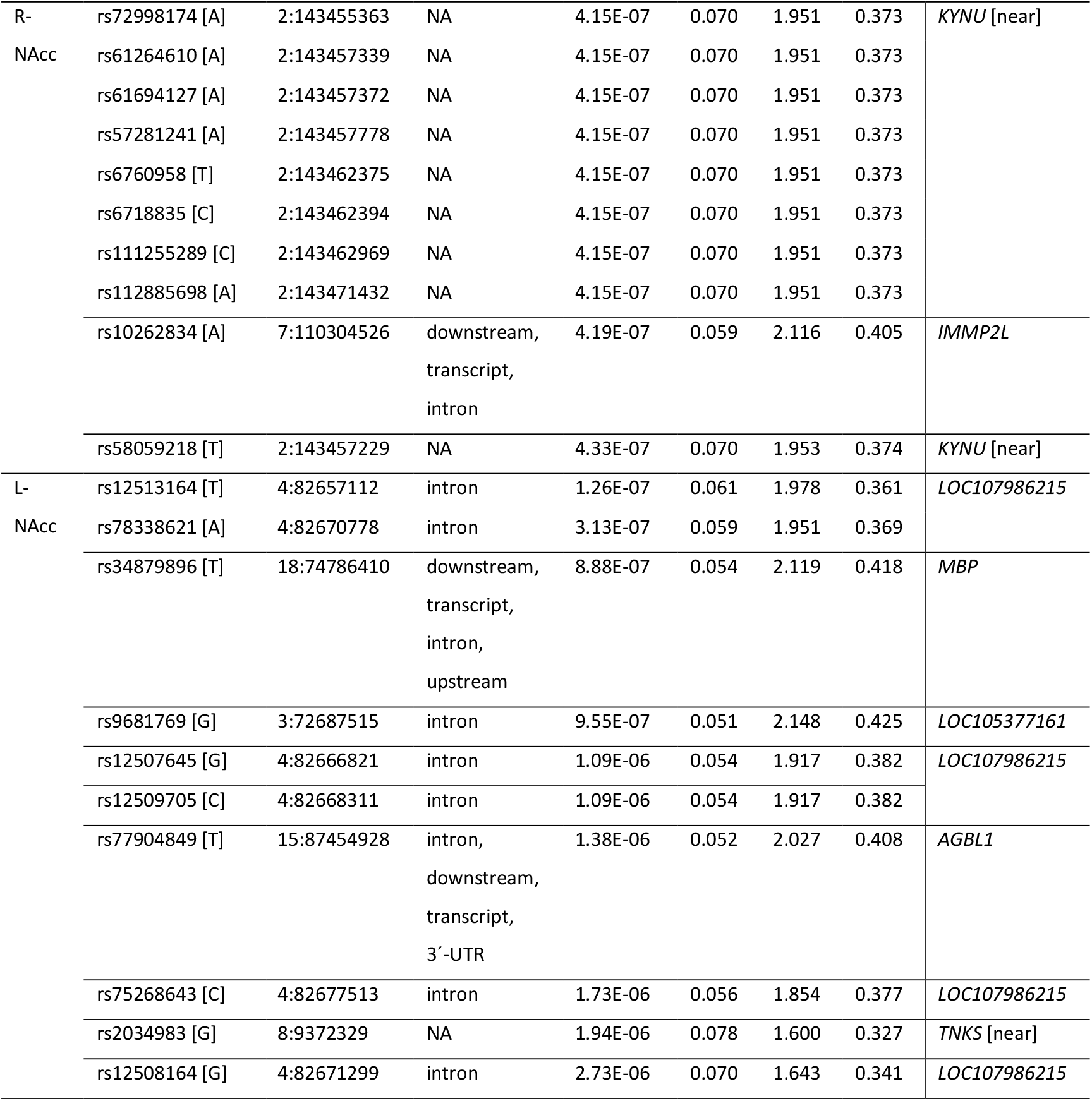
Ten most significant association results from single marker association analyses related to *a priori* ROIs. L: left; R: right; VTA: ventral tegmental area; NAcc: nucleus accumbens; SNP: rs-identifier with counted allele in regression in square brackets; Chromosome: location on chromosome in base pairs according to hg19; SNP type: functional consequence; *P*-value: asymptotic p-value; Freq: allele frequency; Beta: effect size; SE: standard error; Locus: gene annotation according to dbSNP (near gene indicated in square brackets); NA: not available.

Of the ten genetic variants with the lowest p-values associated with activation of the right VTA, one variant was located in *DNAJC13* (p = 2.50E-07). Nine variants were situated in the *ACAD11* (Acyl-CoA Dehydrogenase Family Member 11) and the *NPHP3-ACAD11* (Nephrocystin 3 - Acyl-CoA Dehydrogenase Family Member 11 readthrough transcript) gene, with p-values ranging from p = 4.33E-08 to p = 3.86E-07.

Among the top ten variants associated with the strength of activation in the left VTA, one was located in *SV2C*, three in *FRMD4A* (FERM Domain Containing 4A) and *LOC105376426*, one in *WDR7-OT1* (WDR7 Overlapping Transcript 1), one in *NBEA* (Neurobeachin), and three in *ACAD11* and *NPHP3-ACAD11*, respectively. One variant was assigned to *ACAD11, NPHP3-ACAD11* and *ACKR4* (Atypical Chemokine Receptor 4). Totally, the p-values of these variants ranged from p = 1.51E-06 to p = 3.83E-06.

For the reactivity of the right NAcc, one of the ten variants with the best p-values was located in *IMMP2L*, whereas nine others were situated near the gene *KYNU* (Kynureinase), with overall p-values between p = 4.15E-07 and p = 4.33E-07.

Six of the top ten variants for the strength of association with responses in the left NAcc were located in *LOC107986215*, one in *MBP* (Myelin Basic Protein), one in *LOC105377161* and one in *AGBL1* (ATP/GTP Binding Protein Like 1). One variant was situated near the gene *TNKS* (Tankyrase). P-values of these variants ranged from p = 1.26E-07 to p = 2.73E-06.

To exclude potential biases, we repeated the association analysis of the top ten variants in each brain region from Table 1 including the first and second principal components (PC1 & PC2) as covariates (Supplementary Methods). Repetition including PC1 & PC2 showed that the findings (top hits) were robust (Supplementary Table 2).

### Genetic Association Analyses of Haplotypes

In the haplotype analyses that were performed to strengthen the evidence for the most plausible genes, association signals of gene-spanning haplotypes were revealed only for the *DNAJC13* locus. As described in section 2.7.2, we minimized the number of markers at the *DNAJC13* locus to a haplotype length of five. These five-marker haplotypes were built from the alleles of rs6439356 (A/G, first base position in the haplotype), rs3860499 (A/G, second base position in the haplotype), rs7622472 (A/G, third base position in the haplotype), rs6439360 (A/G, fourth base position in the haplotype) and rs12639254 (A/G, fifth base position in the haplotype).

This five-marker haplotype covers more than 125 kb. In the haplotype-based association analysis, nine different haplotypes were detected. We refer to these five-marker haplotypes as HAP1 to HAP9, i.e. AGAGG (HAP1), AAAGG (HAP2), AGAGA (HAP3), AAAGA (HAP4), GAAGA (HAP5), AGAAA (HAP6), GAAAA (HAP7), AGGAA (HAP8) and GAGAA (HAP9). For more details see Materials and Methods section 2.7.2. For increased response to conditioned stimuli in the right VTA, HAP1 displayed a value of p = 3.21E-007. For more details on all nine haplotypes see Supplementary Table 3.

### Additional Whole-Brain Neuroimaging Analyses for *DNAJC13* Single Marker rs113408797

In order to further explore the effects of *DNAJC13* variant rs113408797 on other brain regions, particularly within the extended reward system (i.e., beside effects on our primary seed regions), additional whole-brain group analyses were conducted comparing groups differing with respect to the SNP (contrast: [heterozygous allele carrier + homozygous minor allele carrier] > homozygous major allele carrier). This analysis demonstrated that, in addition to the effect on reward-related activation in the right VTA (Figure 1), activation in further brain regions of the extended reward system, e.g., in the left vStr (cf. Figure 1) and bilaterally in the anterior insula, was also significantly increased in carriers of the rs113408797 minor allele. Additional group differences depending on the genotype were also detected in regions outside the reward system (see Table 2).

**Figure 1.**
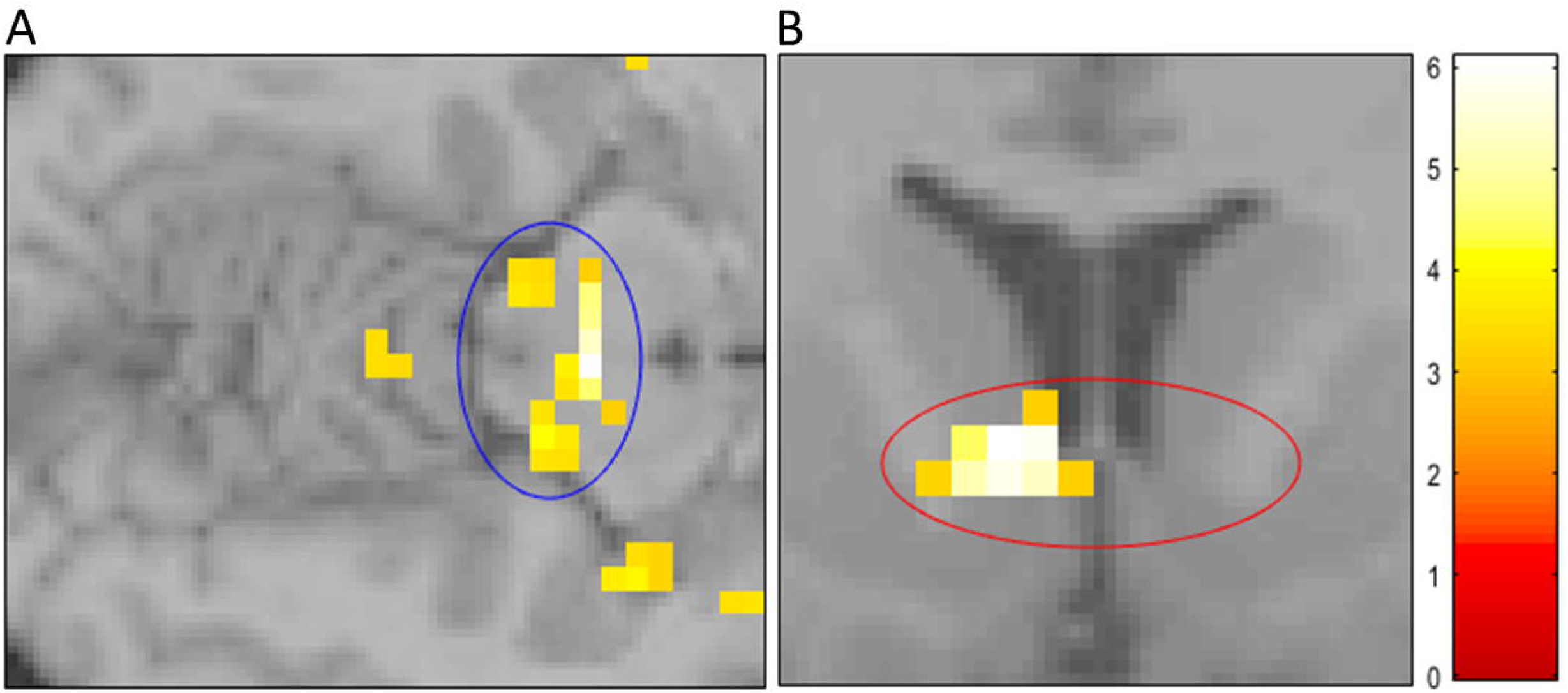
Effects of *DNAJC13* rs113408797 on reward-related activation of the ventral tegmental area (A) and the left ventral striatum (B). Minor allele carriers show increased responses as compared to major allele homozygotes to conditioned reward stimuli in the desire context. For illustration purposes, the T-map of these effects was thresholded at p < 0.001 (see Table 2 for p-values after FWE-correction for the whole brain).

**Table 2.** Effects of *DNAJC13* variant rs113408797 on reward-related brain activation (contrast rs113408797 minor allele carriers > major allele homozygotes; p-values correspond to FWE-correction for the whole brain. L: left; R: right).

| *Brain region* | *MNI coordinates* | *t-value* | *P-value*<sub>FWE-corr.</sub> |
| --- | --- | --- | --- |
| R - ventral tegmental area | 3 -21 -12 | 6.09 | **< 0.001** |
| L - ventral striatum | -6 12 0 | 6.09 | **< 0.001** |
| L - superior frontal gyrus | -12 51 39 | 5.76 | **0.001** |
| frontomedian cortex | -3 33 57 | 4.84 | **0.047** |
| L - anterior insula | -36 9 -6 | 5.49 | **0.003** |
| L - posterior middle temporal gyrus | -51 -42 -3 | 5.49 | **0.003** |
| L - Heschl's transverse convolutions (gyri temporales transversi) | -54 -12 3 | 5.41 | **0.004** |
| L - medial segment of postcentral gyrus | -3 -39 63 | 5.23 | **0.010** |
| L - nucleus reticularis thalami/nucleus caudatus | -9 -9 18 | 5.05 | **0.020** |
| R - anterior insula/deep frontal operculum | 33 30 0 | 4.99 | **0.026** |
| L - posterior superior temporal gyrus/temporoparietal junction cortex | -57 -48 12 | 4.98 | **0.027** |
| L - inferior frontal junction | -36 15 30 | 4.86 | **0.043** |
| L - frontoparietal operculum | -42 -9 21 | 4.84 | **0.047** |
| L - posterior insula | -42 -6 -6 | 4.79 | 0.056 |
| R - inferior frontal gyrus pars triangularis | 48 36 6 | 4.70 | 0.079 |
| R - superior occipital gyrus | 27 -78 39 | 4.68 | 0.084 |
| L - hippocampus | -33 -12 -24 | 4.67 | 0.087 |
| R - gyrus fusiformis | 27 -48 0 | 4.63 | 0.101 |
| L - frontomedian cortex | -6 18 48 | 4.52 | 0.149 |
| L - Broca's area | -57 12 18 | 4.49 | 0.165 |

## DISCUSSION

The objective of our study was to analyze allele-related variation associated with the reactivity of subcortical key regions of the mesolimbic reward system in response to reward stimuli on a genome-wide scale. Consistent with the assumption of polygenic influences on this complex trait, single-marker association analyses revealed several loci scattered across the genome that displayed genetic effects on reward-related brain activation. The most robust finding was obtained in minor allele carriers of *DNAJC13* variant rs113408797 showing increased responses of the VTA and NAcc to conditioned reward stimuli. We also demonstrate a *DNAJC13* haplotype association with reward-related activation in VTA which provides further confirmation for the effects of *DNAJC13* seen in single-marker analysis across the entire gene. This finding may either indicate that rs113408797 constitutes a causal variant and the five markers of the haplotype tag its alleles, or alternatively that all six variants or a combination thereof may each functionally contribute to the effect. On the other hand, possibly all six variants may be markers for a so far undetected causal genetic variation.

The observed influences of *DNAJC13* on the reactivity of the DA system in response to conditioned reward stimuli may be mediated via various molecular mechanisms. While the precise mechanism of action of DNAJC13 on cellular processes is not fully known, some aspects have been described and can therefore be specified in more detail. Particularly, three partly interrelated molecular mechanisms plausibly link the gene product of *DNAJC13* (alias RME-8) to dopaminergic signaling and thereby to reward system functioning (i-iii).

i. DNAJC13 as co-chaperone Due to its function in protein folding, DNAJC13 is involved in synapse maintenance. Without chaperone activity at synapses, deficits in neurotransmission, dismantling of synapses, and degeneration of neurons occur. The function of co-chaperones is to assist heat shock proteins (chaperones) in the cellular process of protein folding and regulate distinct steps of the synaptic vesicle cycle [20]. Specifically, DNAJ-proteins help heat shock protein 70 (Hsp70) to catalyze ATP hydrolysis and to recruit other chaperones that cause a conformational change of unfolded polypeptide chains and release proteins after they have been properly folded [21]. With respect to the reward system, Hsp70 has been shown to protect dopaminergic neurons from toxic influences in models of Parkinson’s disease, both *in vitro* in primary mesencephalic neurons and *in vivo* in the substantia nigra pars compacta [22]. Therefore, quantitative or qualitative changes in DNAJC13 may induce differences in folding and protein aggregation via Hsp70, which is relevant for dopaminergic signaling.
ii. DNAJC13 on vesicle dynamics DNAJC13 influences coated vesicle endocytosis via clathrin, which plays an important role in the formation of coated vesicles. This is of high relevance to dopaminergic signaling, because the internalization of dopaminergic D2 receptors in the prefrontal cortex is a clathrin-dependent process in primates [23]. Mechanistically, DNAJC13 was shown to interact with Hsp70 and to use a phosphoinositide-binding motive to regulate the endosomal clathrin dynamics [24,25]. With respect to neuropsychiatric disorders, many neurobiological aspects of psychoses are clathrin-dependent, e.g., synaptic dysfunction, alterations in white matter and aberrant neurodevelopment. Furthermore, as many antipsychotics affect clathrin-interacting proteins, it has been proposed that changes induced in the interactome of clathrin may offer potential treatments [26].
iii. DNAJC13 in autophagy DNAJC13 plays a role in the stabilization of cellular protein homeostasis. A disruption of this process results in the accumulation of protein aggregates, which is known to contribute to neurodegenerative diseases [27]. Furthermore, DNAJC13 influences protein degradation via the autophagy pathway, whereby the cell removes misfolded proteins due to an association with the retromer and the WASH complex. The retromer ensures degradation of autophagic cargo by maintaining lysosome function [28], whereas the WASH complex is required for lysosomal maturation and recycling, and therefore for efficient autophagic and phagocytic digestion [29]. Comprehensive knockdown and overexpression experiments of *DNAJC13* clarified that it acts as a positive modulator in autophagic processes [30].

Although our study clearly suggests that genetic variation of the *DNAJC13* locus impacts on mesolimbic reward system responses to conditioned reward stimuli, currently it cannot be decided which of these molecular mechanisms, or rather a combination of several options, mediates these effects.

So far, most reported genetic associations with reward system activity are based on core dopaminergic candidate genes where the biological mechanism of action is obvious based on previous biochemical, cellular and animal work. The effects of such candidate genes on neurofunctional mechanisms have also been investigated using functional neuroimaging in humans. For instance, firing of dopaminergic neurons in the VTA, which form synapses with neurons in the NAcc, were recognized as particularly important for the encoding of reward information [31]. After the release of DA, the action of the neurotransmitter is terminated by re-uptake into the presynapse of the VTA neuron through the DA transporter (*DAT1*). Regarding *DAT1*, genetic variation is already known to influence reward system activity [32,33]. At the postsynapse, which belongs to the neuron in the NAcc, DA binds to receptors. Based on this receptor activation a signaling cascade involving a chain of intracellular reactions on transcription factors, e.g., cAMP response element-binding protein 1 (*CREB1*), is initiated. *CREB1* in turn influences gene expression and the cellular response. Genetic variation of *CREB1* was also shown to affect reward system functioning in a previous fMRI study [34].

Some other genes whose relationship to dopaminergic neurotransmission is less obvious, have also been reported to influence activity within the extended reward system. For example, this is the case for genetic variation in mitotic arrest deficient 1 like 1 (*MAD1L1*), whose primary role is cell cycle control. *MAD1L1* has initially been identified as a potential susceptibility factor for bipolar disorder in a GWAS [35]. Subsequently, by imaging genetics analyses, influences of *MAD1L1* on reward system reactivity were shown. Healthy risk allele carriers showed a significant deficit of their bottom-up response to conditioned reward stimuli in the bilateral VTA and vStr [36]. Another gene that is indirectly linked to dopaminergic neurotransmission is vacuolar protein sorting-associated protein 4a (*VPS4A*) [16]. The genetic variation of this gene was implicated in a GWAS for reward anticipation. The gene product of *VPS4A* functions in intracellular protein transport and was suggested to affect catecholaminergic (dopaminergic and/or adrenergic) activity and reward processing [16].

In a similar way, considering that *DNAJC13* is involved in Parkinson’s disease [37-39] and schizophrenia [40,41], which are both related to dopaminergic dysfunctions, indirect evidence also links this molecule to dopamine. The exact mechanisms how *DNAJC13* may act in dopamine system functioning (and dysfunction) still have to be elucidated in future research.

For example, one may hypothesize that in *DNAJC13* allele and/or haplotype carriers possible alterations in reward system functioning may be represented by subthreshold symptoms and, in particular, may contribute to clinical disease manifestation according to the concept of an endophenotype. The mesolimbic reward system works in concert with cortical areas to execute reward-related behaviors [7,42]. Our findings provide first empirical evidence that genetic variation in *DNAJC13* may potentially increase individual vulnerability for neurological and psychiatric disorders like Parkinson’s disease and schizophrenia by modulating the functional interplay of brain regions within the extended reward system. Thus, *DNAJC13* could be further investigated for its possible role in pathogenesis of more specific forms of schizophrenia and/or affective disorders that are related to extended reward system dysfunction. If this gene shows increased effects in those subforms, in which dopaminergic mechanisms play a major role, its future investigation may elucidate the underlying pathomechanisms.

Our study is limited by the fact that the sample size is relatively small for investigating genetic associations at a genome-wide level. On the other hand, careful application of strict inclusion and exclusion criteria secured that the sample is very homogeneous with respect to potentially confounding effects on neurofunctional systems that may result from differences in age, education and task performance levels as well as from pathological changes due to neurological or psychiatric disorders. This may be one important aspect of study design that contributed to the rather large genetic effects observed within this very homogeneous sample. Nevertheless, as usual our findings will have to be replicated in future studies.

The present investigation is one of the first studies that analyzed genetic variation acting on the mesolimbic reward system on a genome-wide scale. In sum, our findings provide direct empirical evidence that *DNAJC13* influences responses of brain regions within the extended reward system to conditioned stimuli. To understand the genetic contribution of *DNAJC13* to neural activation strength in more detail, future studies may examine the links of this gene to other molecular factors that play a role for reward system activation.

## ACKNOWLEDGMENTS

We thank all subjects who participated in this study. We thank Prof. Dr. Dr. Elisabeth Binder and Monika Rex-Haffner, Max-Planck-Institute for Psychiatry, Munich for genome-wide SNP genotyping and Maria Keil, Center for Translational Research in Systems Neuroscience and Psychiatry, Department of Psychiatry and Psychotherapy, Georg-August-University Göttingen, Göttingen, for support in recruitment and fMRI investigation of volunteers.

## AUTHOR CONTRIBUTIONS

J.T., K.E.E., B.K., S.A., S.R. and O.G. made substantial contributions to the acquisition, analysis and/or interpretation of the data. J.T., K.E.E. and O.G. contributed by drafting the manuscript and/or revising it critically for important intellectual content. O.G. conceptualized and designed the study.

## SUPPLEMENTARY INFORMATION

Supplementary information accompanies the paper.

## REFERENCES

1 Gottesman, II, Gould TD. The endophenotype concept in psychiatry: etymology and strategic intentions. Am J Psychiatry. 2003;160(4):636–45.

2 Cannon TD, Keller MC. Endophenotypes in the genetic analyses of mental disorders. Annu Rev Clin Psychol. 2006;2:267–90.

3 Pearlson GD, Calhoun VD. Convergent approaches for defining functional imaging endophenotypes in schizophrenia. Front Hum Neurosci. 2009;3:37.

4 Grimm O, Heinz A, Walter H, Kirsch P, Erk S, Haddad L, et al. Striatal response to reward anticipation: evidence for a systems-level intermediate phenotype for schizophrenia. JAMA Psychiatry. 2014;71(5):531–9.

5 Schultz W, Dayan P, Montague PR. A neural substrate of prediction and reward. Science. 1997;275(5306):1593–9.

6 Trost S, Diekhof EK, Zvonik K, Lewandowski M, Usher J, Keil M, et al. Disturbed anterior prefrontal control of the mesolimbic reward system and increased impulsivity in bipolar disorder. Neuropsychopharmacology. 2014;39(8):1914–23.

7 Diekhof EK, Gruber O. When desire collides with reason: functional interactions between anteroventral prefrontal cortex and nucleus accumbens underlie the human ability to resist impulsive desires. J Neurosci. 2010;30(4):1488–93.

8 Juckel G, Schlagenhauf F, Koslowski M, Filonov D, Wustenberg T, Villringer A, et al. Dysfunction of ventral striatal reward prediction in schizophrenic patients treated with typical, not atypical, neuroleptics. Psychopharmacology (Berl). 2006;187(2):222–8.

9 Subramaniam K, Hooker CI, Biagianti B, Fisher M, Nagarajan S, Vinogradov S. Neural signal during immediate reward anticipation in schizophrenia: Relationship to real-world motivation and function. Neuroimage Clin. 2015;9:153–63.

10 Richter A, Petrovic A, Diekhof EK, Trost S, Wolter S, Gruber O. Hyperresponsivity and impaired prefrontal control of the mesolimbic reward system in schizophrenia. J Psychiatr Res. 2015;71:8–15.

11 Gradin VB, Waiter G, O’Connor A, Romaniuk L, Stickle C, Matthews K, et al. Salience network-midbrain dysconnectivity and blunted reward signals in schizophrenia. Psychiatry Res. 2013;211(2):104–11.

12 Diekhof EK, Keil M, Obst KU, Henseler I, Dechent P, Falkai P, et al. A functional neuroimaging study assessing gender differences in the neural mechanisms underlying the ability to resist impulsive desires. Brain Res. 2012;1473:63–77.

13 Brett M, Anton J-L, Valabregue R, Poline J-B. Region of Interest Analysis Using an SPM Toolbox [Abstract]. Neuroimage. 2002;16.

14 dbSNP_database. https://www.ncbi.nlm.nih.gov/snp/. Retrieved: June 2020.

15 Stacey D, Bilbao A, Maroteaux M, Jia T, Easton AC, Longueville S, et al. RASGRF2 regulates alcohol-induced reinforcement by influencing mesolimbic dopamine neuron activity and dopamine release. Proc Natl Acad Sci U S A. 2012;109(51):21128–33.

16 Jia T, Macare C, Desrivieres S, Gonzalez DA, Tao C, Ji X, et al. Neural basis of reward anticipation and its genetic determinants. Proc Natl Acad Sci U S A. 2016;113(14):3879–84.

17 Barrett JC, Fry B, Maller J, Daly MJ. Haploview: analysis and visualization of LD and haplotype maps. Bioinformatics. 2005;21(2):263–5.

18 Gabriel SB, Schaffner SF, Nguyen H, Moore JM, Roy J, Blumenstiel B, et al. The structure of haplotype blocks in the human genome. Science. 2002;296(5576):2225–9.

19 Purcell S, Neale B, Todd-Brown K, Thomas L, Ferreira MA, Bender D, et al. PLINK: a tool set for whole-genome association and population-based linkage analyses. Am J Hum Genet. 2007;81(3):559–75.

20 Gorenberg EL, Chandra SS. The Role of Co-chaperones in Synaptic Proteostasis and Neurodegenerative Disease. Front Neurosci. 2017;11:248.

21 Langer T, Lu C, Echols H, Flanagan J, Hayer MK, Hartl FU. Successive action of DnaK, DnaJ and GroEL along the pathway of chaperone-mediated protein folding. Nature. 1992;356(6371):683–9.

22 Nagel F, Falkenburger BH, Tonges L, Kowsky S, Poppelmeyer C, Schulz JB, et al. Tat-Hsp70 protects dopaminergic neurons in midbrain cultures and in the substantia nigra in models of Parkinson’s disease. J Neurochem. 2008;105(3):853–64.

23 Paspalas CD, Rakic P, Goldman-Rakic PS. Internalization of D2 dopamine receptors is clathrin-dependent and select to dendro-axonic appositions in primate prefrontal cortex. Eur J Neurosci. 2006;24(5):1395–403.

24 Chang HC, Hull M, Mellman I. The J-domain protein Rme-8 interacts with Hsc70 to control clathrin-dependent endocytosis in Drosophila. J Cell Biol. 2004;164(7):1055–64.

25 Xhabija B, Vacratsis PO. Receptor-mediated Endocytosis 8 Utilizes an N-terminal Phosphoinositide-binding Motif to Regulate Endosomal Clathrin Dynamics. J Biol Chem. 2015;290(35):21676–89.

26 Schubert KO, Focking M, Prehn JH, Cotter DR. Hypothesis review: are clathrin-mediated endocytosis and clathrin-dependent membrane and protein trafficking core pathophysiological processes in schizophrenia and bipolar disorder? Mol Psychiatry. 2012;17(7):669–81.

27 Klaips CL, Jayaraj GG, Hartl FU. Pathways of cellular proteostasis in aging and disease. J Cell Biol. 2018;217(1):51–63.

28 Maruzs T, Lorincz P, Szatmari Z, Szeplaki S, Sandor Z, Lakatos Z, et al. Retromer Ensures the Degradation of Autophagic Cargo by Maintaining Lysosome Function in Drosophila. Traffic. 2015;16(10):1088–107.

29 King JS, Gueho A, Hagedorn M, Gopaldass N, Leuba F, Soldati T, et al. WASH is required for lysosomal recycling and efficient autophagic and phagocytic digestion. Mol Biol Cell. 2013;24(17):2714–26.

30 Besemer AS, Maus J, Ax MDA, Stein A, Vo S, Freese C, et al. Receptor-mediated endocytosis 8 (RME-8)/DNAJC13 is a novel positive modulator of autophagy and stabilizes cellular protein homeostasis. Cell Mol Life Sci. 2021;78(2):645–60.

31 Dichter GS, Damiano CA, Allen JA. Reward circuitry dysfunction in psychiatric and neurodevelopmental disorders and genetic syndromes: animal models and clinical findings. J Neurodev Disord. 2012;4(1):19.

32 Dreher JC, Kohn P, Kolachana B, Weinberger DR, Berman KF. Variation in dopamine genes influences responsivity of the human reward system. Proc Natl Acad Sci U S A. 2009;106(2):617–22.

33 Hahn T, Heinzel S, Dresler T, Plichta MM, Renner TJ, Markulin F, et al. Association between reward-related activation in the ventral striatum and trait reward sensitivity is moderated by dopamine transporter genotype. Hum Brain Mapp. 2011;32(10):1557–65.

34 Wolf C, Mohr H, Diekhof EK, Vieker H, Goya-Maldonado R, Trost S, et al. CREB1 Genotype Modulates Adaptive Reward-Based Decisions in Humans. Cereb Cortex. 2016;26(7):2970–81.

35 Cichon S, Muhleisen TW, Degenhardt FA, Mattheisen M, Miro X, Strohmaier J, et al. Genome-wide association study identifies genetic variation in neurocan as a susceptibility factor for bipolar disorder. Am J Hum Genet. 2011;88(3):372–81.

36 Trost S, Diekhof EK, Mohr H, Vieker H, Kramer B, Wolf C, et al. Investigating the Impact of a Genome-Wide Supported Bipolar Risk Variant of MAD1L1 on the Human Reward System. Neuropsychopharmacology. 2016;41(11):2679–87.

37 Appel-Cresswell S, Rajput AH, Sossi V, Thompson C, Silva V, McKenzie J, et al. Clinical, positron emission tomography, and pathological studies of DNAJC13 p.N855S Parkinsonism. Mov Disord. 2014;29(13):1684–7.

38 Vilarino-Guell C, Rajput A, Milnerwood AJ, Shah B, Szu-Tu C, Trinh J, et al. DNAJC13 mutations in Parkinson disease. Hum Mol Genet. 2014;23(7):1794–801.

39 Gustavsson EK, Trinh J, Guella I, Vilarino-Guell C, Appel-Cresswell S, Stoessl AJ, et al. DNAJC13 genetic variants in parkinsonism. Mov Disord. 2015;30(2):273–8.

40 Piras IS, Manchia M, Huentelman MJ, Pinna F, Zai CC, Kennedy JL, et al. Peripheral Biomarkers in Schizophrenia: A Meta-Analysis of Microarray Gene Expression Datasets. Int J Neuropsychopharmacol. 2019;22(3):186–93.

41 Wu Y, Yao YG, Luo XJ. SZDB: A Database for Schizophrenia Genetic Research. Schizophr Bull. 2017;43(2):459–71.

42 Grace AA, Floresco SB, Goto Y, Lodge DJ. Regulation of firing of dopaminergic neurons and control of goal-directed behaviors. Trends Neurosci. 2007;30(5):220–7.

