## Supplement for "*DNAJC13* influences responses of the extended reward system to conditioned stimuli: a genome-wide association study"

**SUPPLEMENTARY METHODS**

1. *Genotypes QC and Imputation*

The genotype quality control was performed using following standard parameters for retaining subjects and SNPs: SNP missingness < 0.05 (before sample removal); subject missingness < 0.02; autosomal heterozygosity deviation (| Fhet | < 0.2); SNP missingness < 0.02 (after sample removal); difference in SNP missingness between cases and controls < 0.02; SNP Hardy-Weinberg equilibrium (p < 1E-06 in controls or p < 1E-10 in cases) and male subjects with heterozygosity rate for chromosome X > 0.5 and females subjects < 0.5. The population outliers were excluded based on principal component analysis (visual inspection).

The genotype imputation was performed using the pre-phasing/imputation stepwise approach implemented in EAGLE/MINIMAC3 (http://genome.sph.umich.edu/wiki/Minimac; MINIMAC, RRID:SCR_009292) (with a variable chunk size of 132 genomic chunks and default parameters). The imputation reference set consisted of 54,330 phased haplotypes with 36,678,882 variants from the publicly available HRC reference (https://ega-archive.org/datasets/EGAD00001002729; European Genome phenome Archive RRID:SCR_004944).

The relatedness was tested using a subset of 80,207 SNPs obtained with a stringent quality criterion (INFO > 0.8, missingness < 1%, minor allele frequency > 0.05) and by applying a LD pruning (*r*2 > 0.02). One member out of the cryptically related pairs (Pi_hat > 0.2) were randomly excluded following the preferential retention of cases over controls.

2. *Quantil-quantil (QQ) plots and genomic inflation factor lambda*

QQ plots from p-values were created using the Manhattan Plotter Software (https://www.biologiaevolutiva.org/~cmorcillo/tools/ManhattanPlotter/ManhattanPlotter.htm), by starting the program in the visual mode, using PLINK v2.00a2.3 (64-bit (24 Jan 2020); https://www.cog-genomics.org/plink/2.0/, PLINK RRID:SCR_001757) output files as ManhattanPlotter input files (column labels SNP, CHR, POS, P), and the Manhattan Plotter menu option “QQ plot”. Genomic inflation factor Lambda was calculated with Plink v1.90b6.9, using the commands --assoc and --adjust. For each of the four brain regions, the lambda value was taken as given in the log file produced by Plink v1.90b6.9 calculation, in the line “--adjust: Genomic inflation est. lambda (based on median chisq)”.

3. *Regional association plots*

Regional association plots were produced using partial PLINK v2.00a2.3 output files, for top chromosomal regions from Table 1 of the main text, with a +/- 250 kb window around relevant variants. The Single Nucleotide Polymorphisms Annotator (Arnold et al., 2015; https://snipa.helmholtz-muenchen.de/snipa/?task=regional_association_plot) was used for plotting the trait-associated variants of each region, and to display the strength of association (-log10(P)). Annotations used by the Single Nucleotide Polymorphisms Annotator were the grch37/1kgpp3v5 variant set (population "eur"), and ensembl87 (for functional annotation).

4. *Allele frequencies*

Allele frequencies of the n = 214 individuals were calculated using the --freq command in PLINK v2.00a2.3 (Supporting Table 1).

5. *Genetic association analysis of the top ten variants including PC1 & PC2 as covariates*

To exclude potential biases, we repeated the analysis of the top ten variants in each brain region with the first two principle components (PC1 & PC2) as covariates. To retrieve association p-values, variants of the genotyped dataset were linkage-disequilibrium pruned using PLINK v2.00a2.3, using --maf 0.05, --geno 0.01, --prune and --indep-pairwise 200 50 0.02 commands, resulting in a file plink2.prune.in that contained 11430 markers. In a second step, for calculation of eigenvalues and eigenvectors with PLINK v2.00a2.3, the commands --extract plink2.prune.in and --pca were used. Subsequently, the first two principle components (PC1 & PC2) of the output file plink2.eigenvec were considered as covariates in association testing in PLINK v2.00a2.3, using the --covar command. All other parameters were chosen as described for the calculation without PCA components in the Material and Methods section of the main text. Repetition of the calculation with PC1 & PC2 showed that findings of top hits (top ten) hold with PCA as covariates (Supporting Table 2).

**SUPPLEMENTARY FIGURES**

Supplementary Figure 1. 2D plot of principal component 1 (PC1) and principal component 2 (PC2) showing the population structure prior to the exclusion of individuals. Individuals of the present study (crosses) are presented in the context of further individuals from related studies (circles). Visible outliers were identified by thresholds of ~ PC1 > 0.03 and PC2 > 0.07.

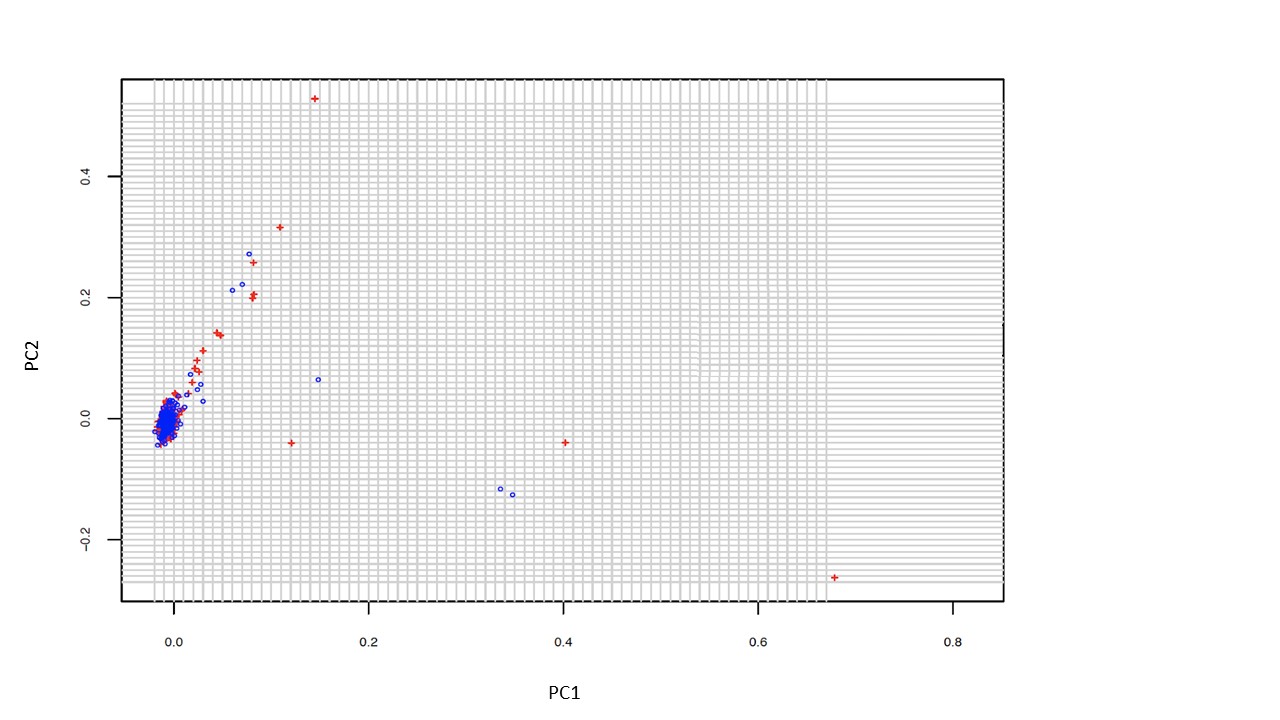

Supplementary Figure 2. Quantil-Quantil (QQ) plots and values for genomic inflation estimates of lambda (based on median chisq).

| a) *L-VTA* (lambda = 1)   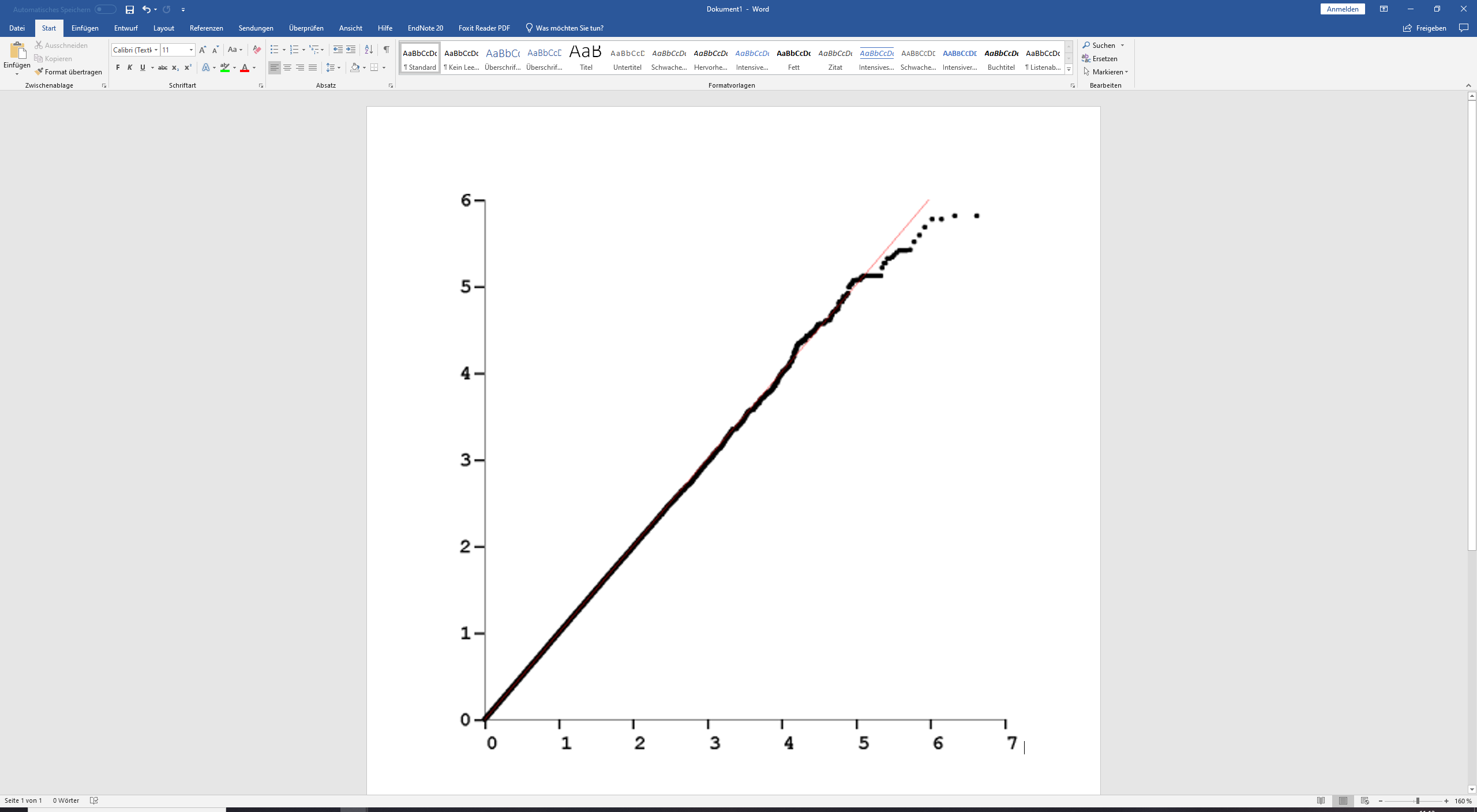 | b) *R-VTA* (lambda = 1)   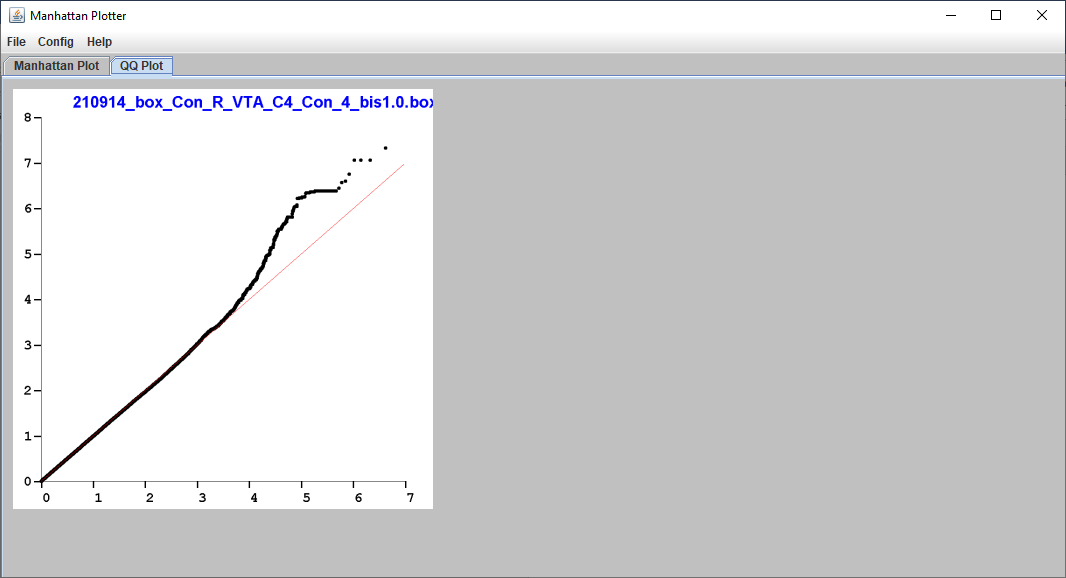 |
| --- | --- |
| c) *L-NAcc* (lambda = 1)   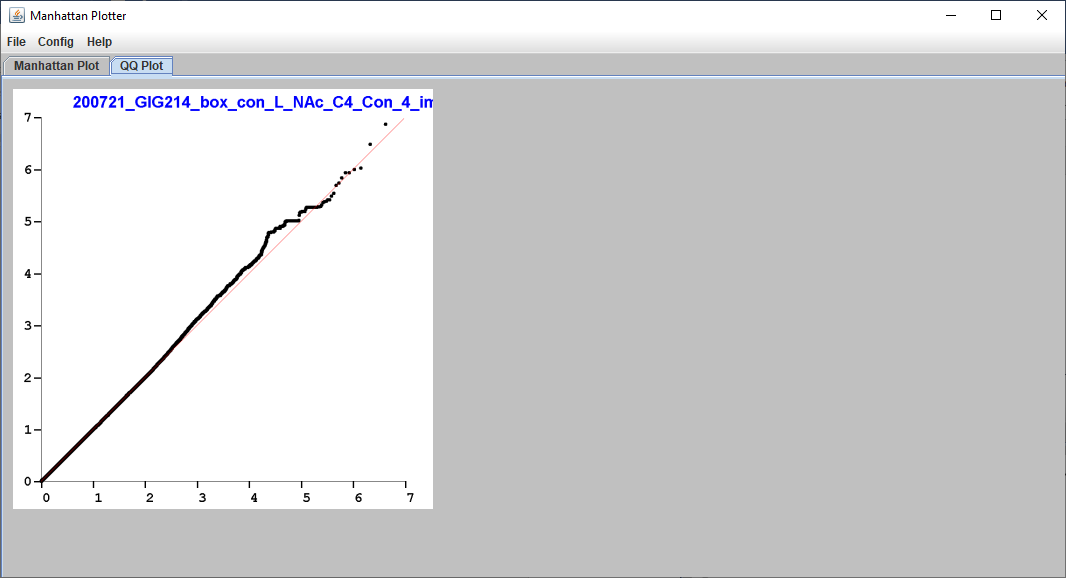 | d) *R-NAcc* (lambda = 1.0048)   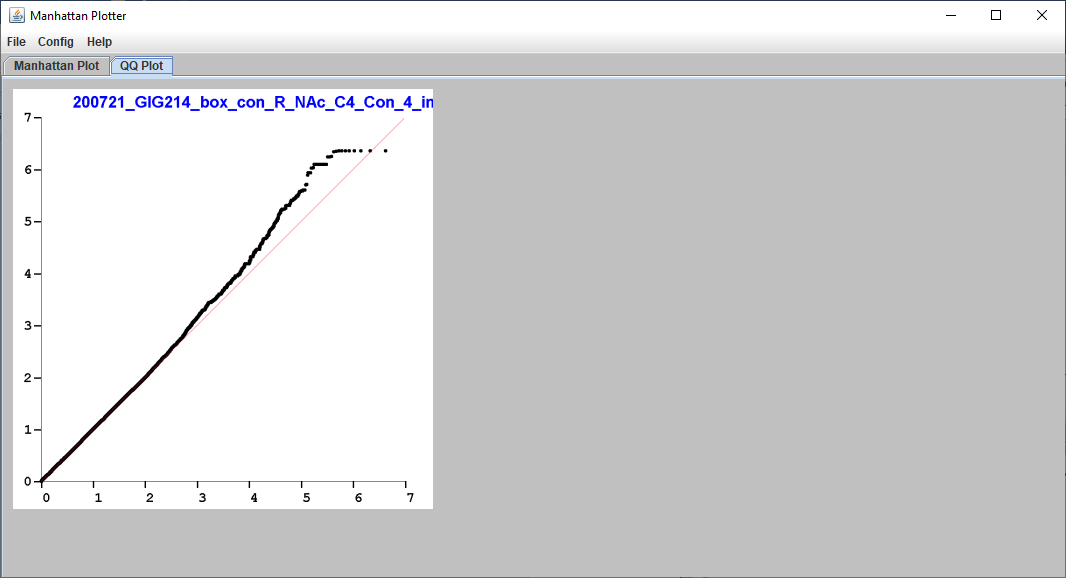 |

Supplementary Figure 3. Regional association plots for the top chromosomal regions from Table 1 of the main text. The colors show, in respect to the top variant of each region, the pairwise LD-correlations of all SNPs of the plots. The light blue lines in the plots display the estimated recombination rates. The dark green arrows at the bottom of the plots show genes and their orientation. The symbols of the variants represent functional regulatory annotations. a) R-VTA; b) L-VTA; c) R-NAcc; d) L-NAcc.

a) **R-VTA**

rs113408797 at chr3q22.1

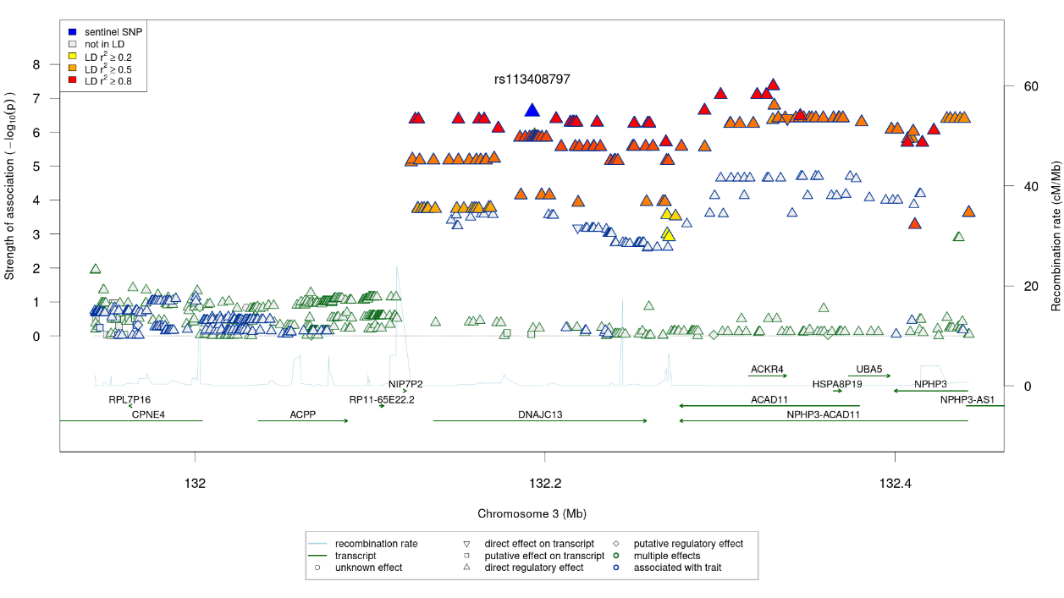

b) **L-VTA**

rs57131074 at chr5q13.3
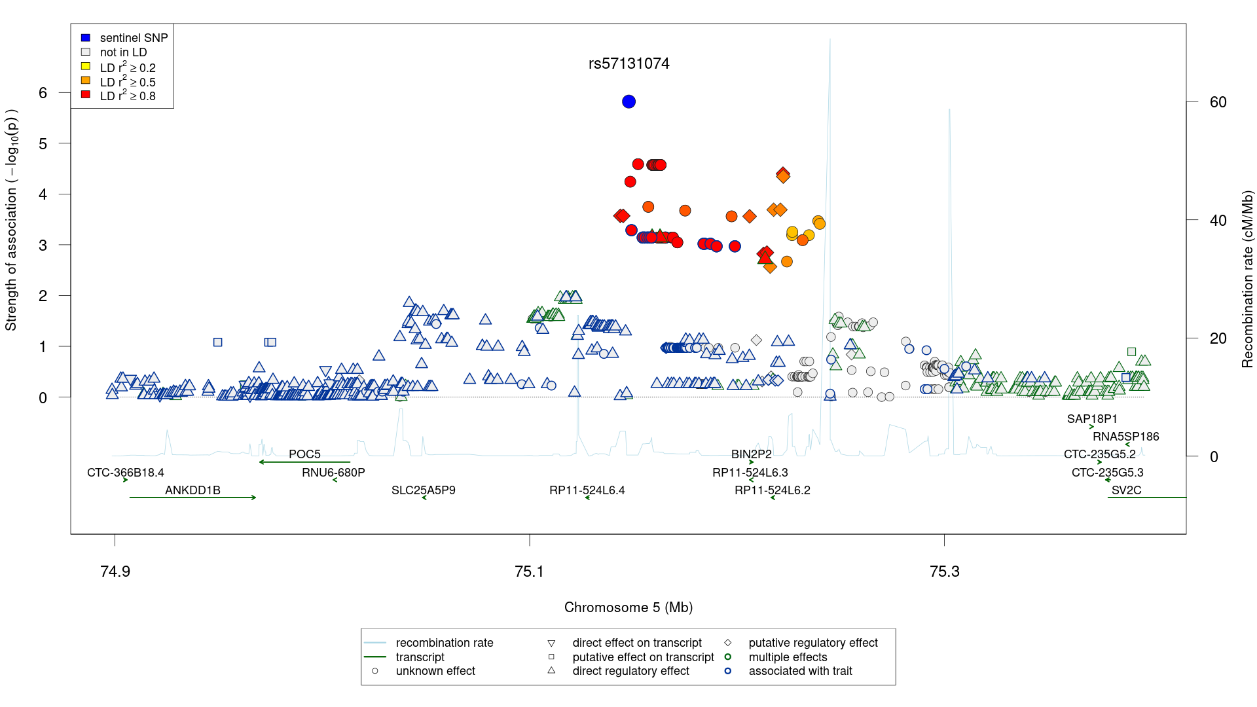

rs10082442 at chr10p13

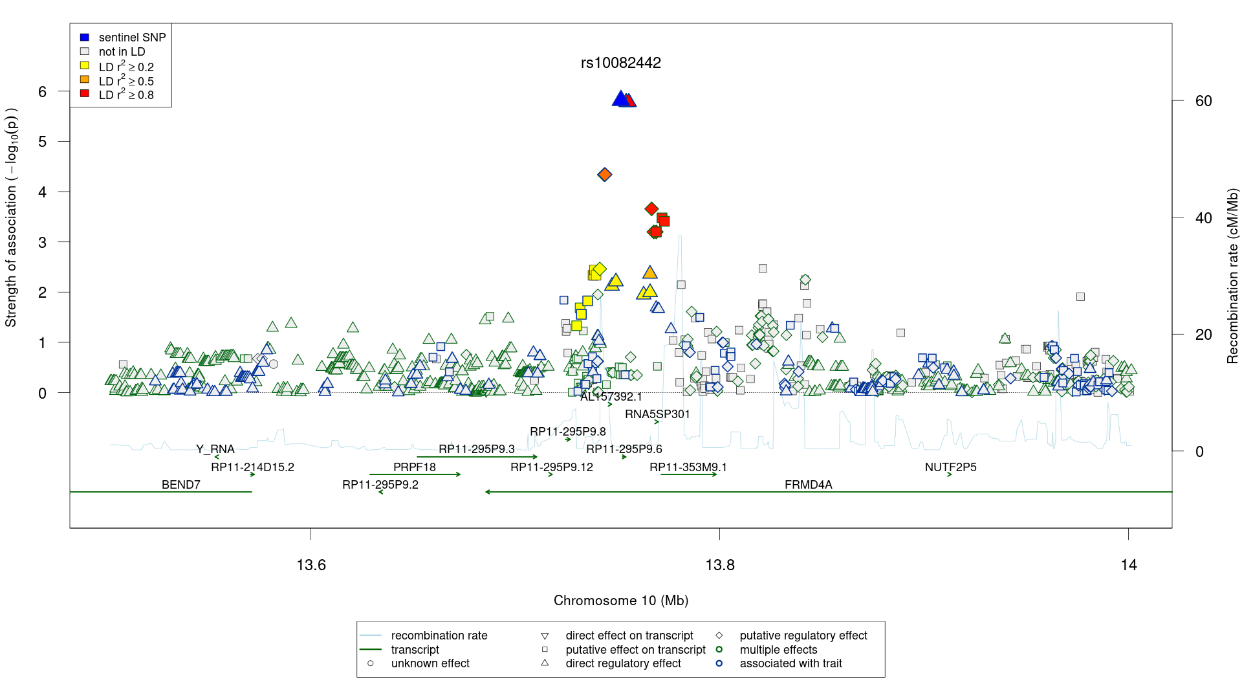

rs1657381 at chr18q21.31

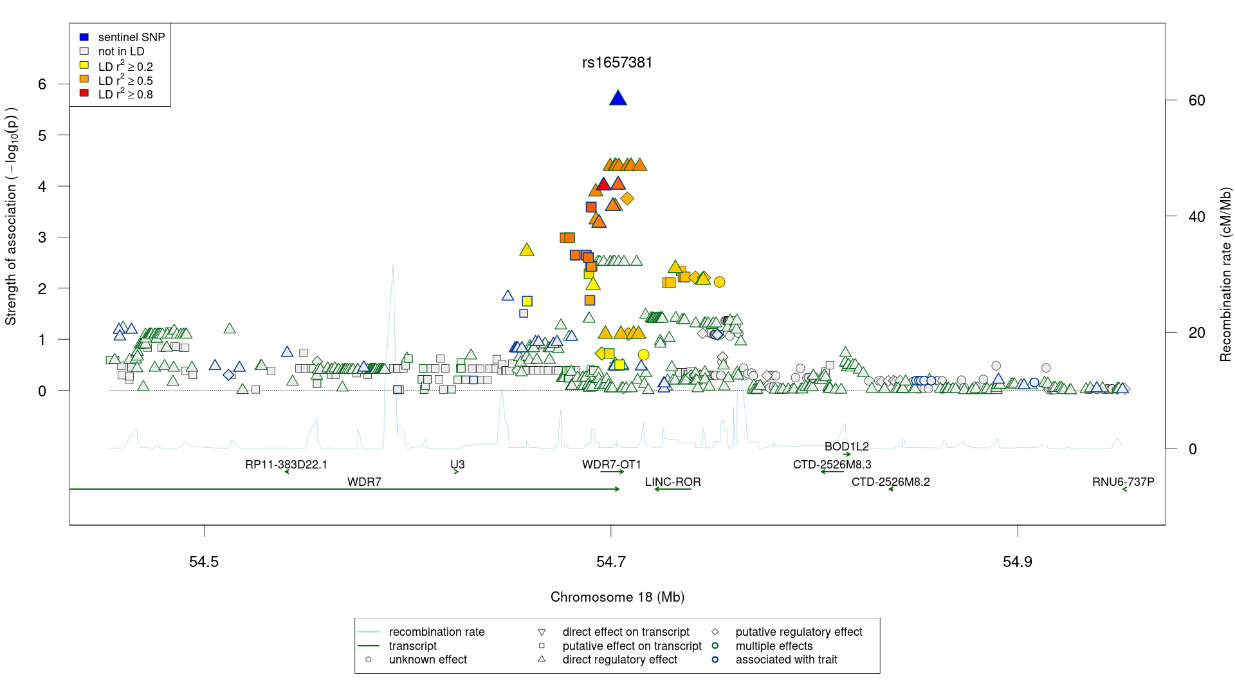

rs1777670 at chr13q13.3

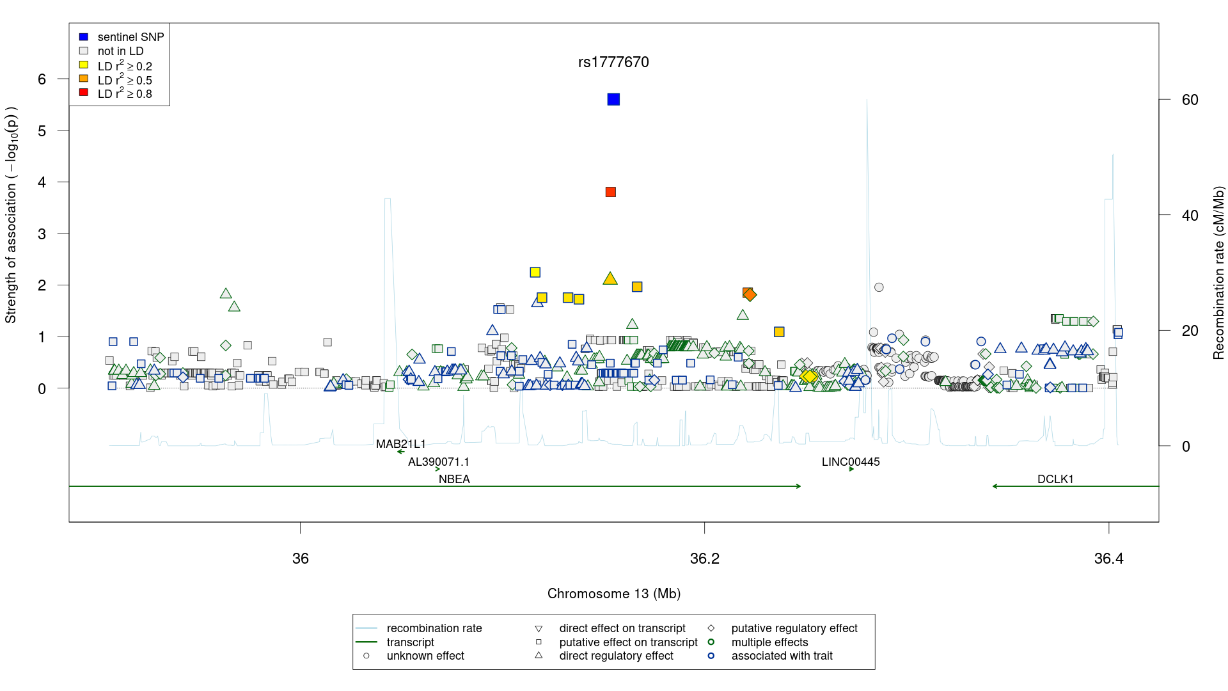

rs141727949 at chr3q22.1

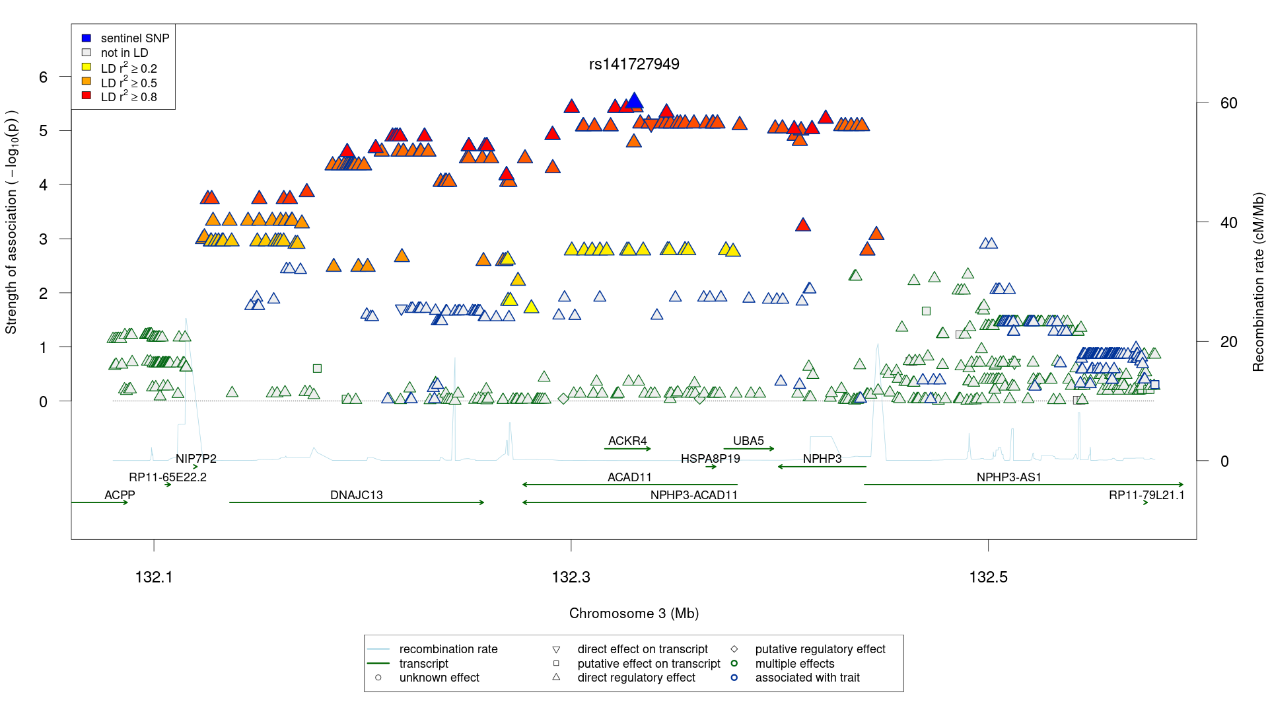

c) **R-NAcc**

rs72998174 at chr2q22.2

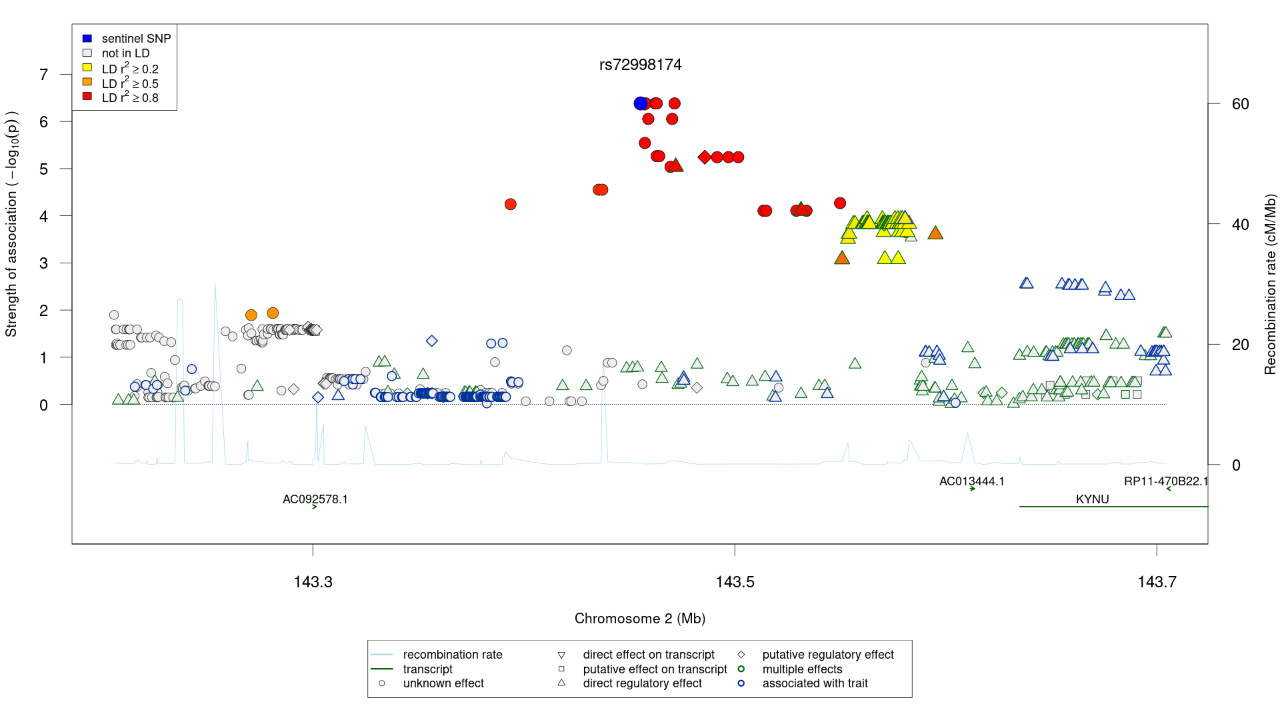

rs10262834 at chr7q31.1

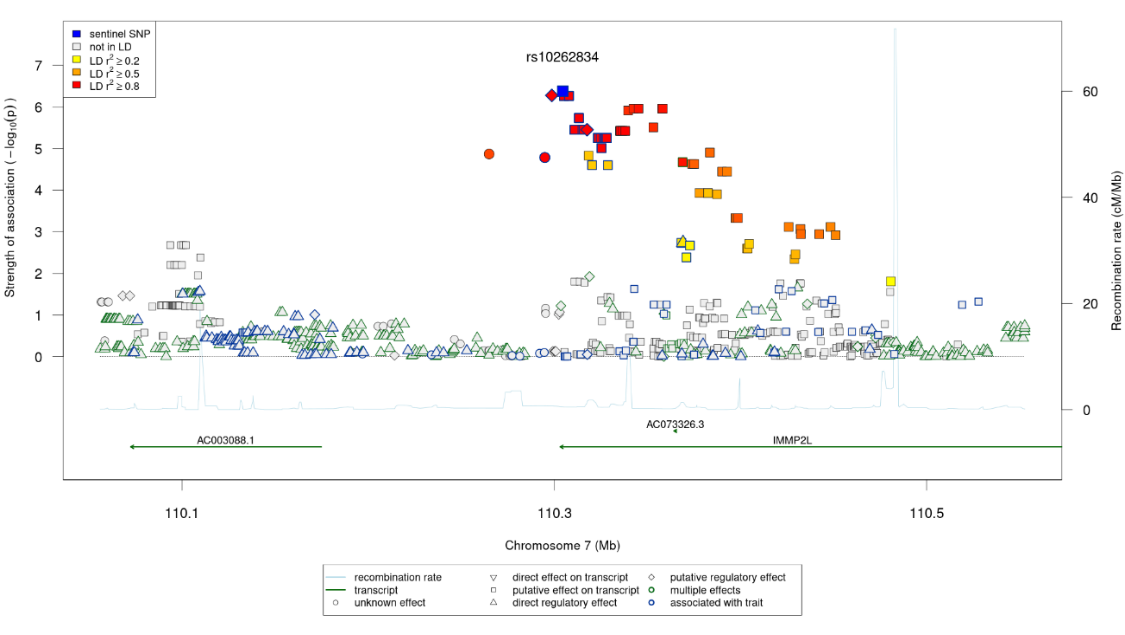

d) **L-NAcc**

rs12513164 at chr4q21.22

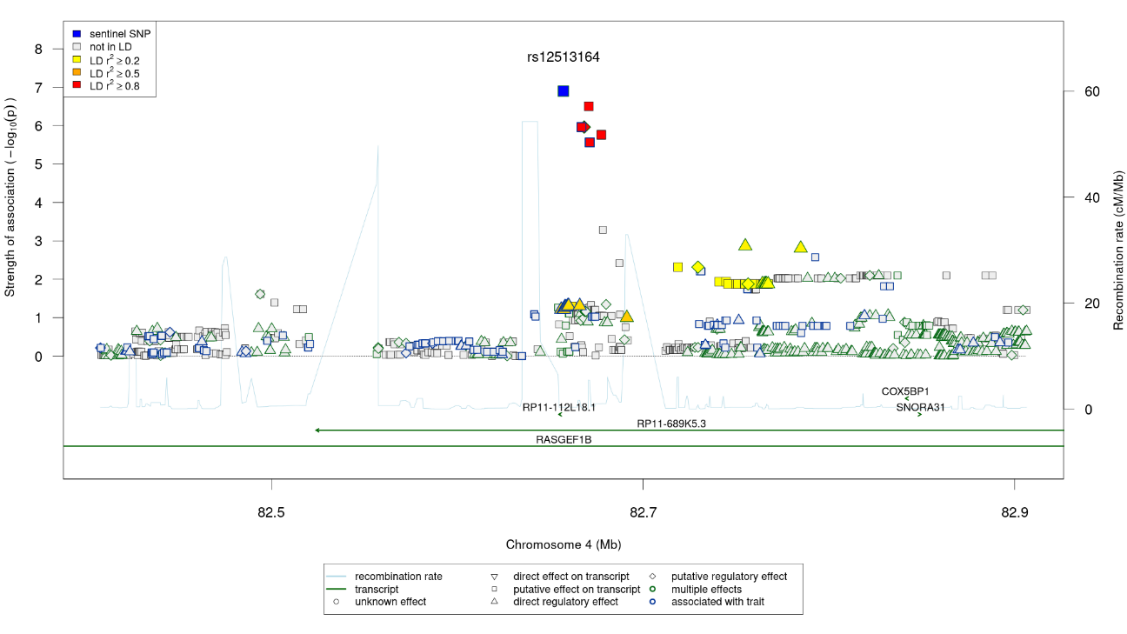

rs34879896 at chr18q23

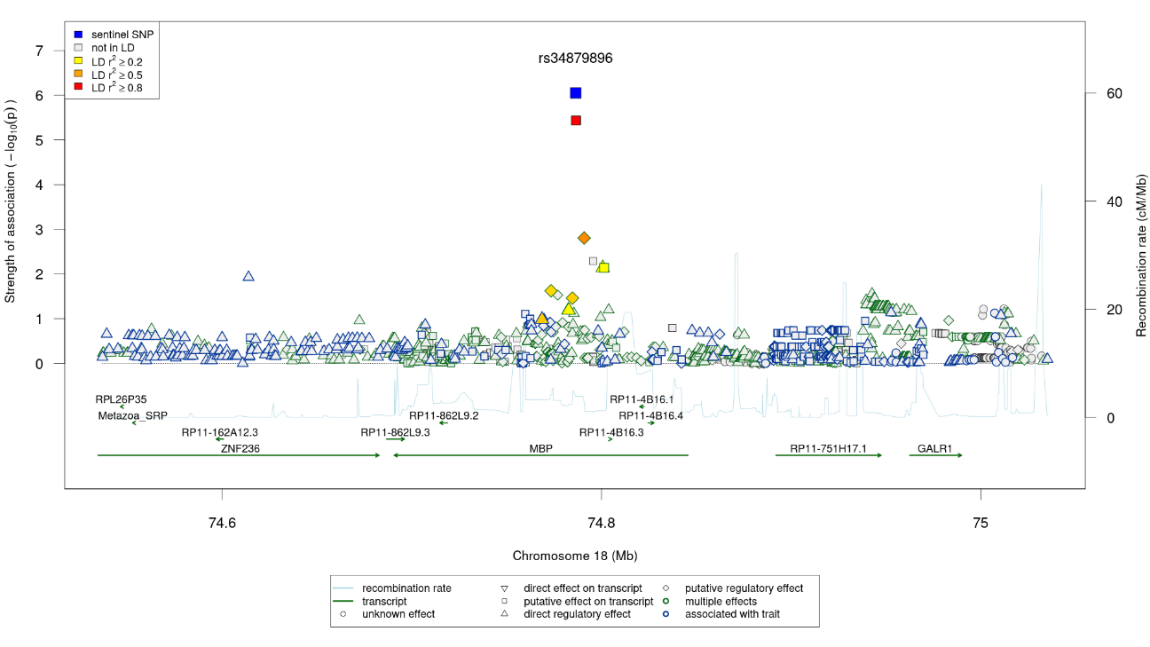

rs9681769 at chr3p13

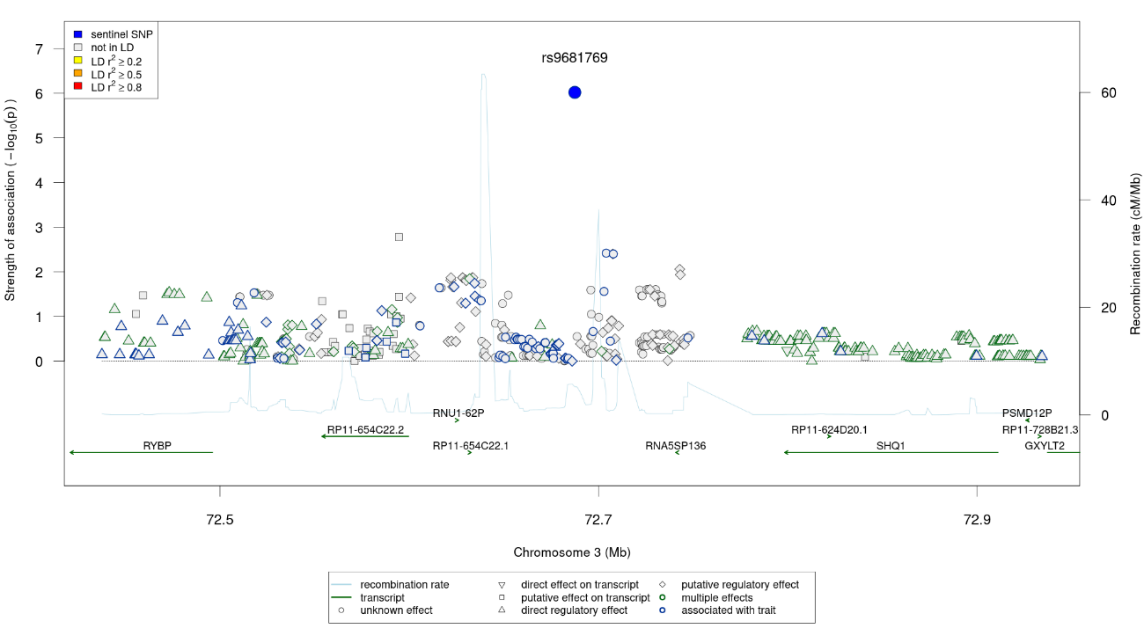

rs77904849 at chr15q25.3

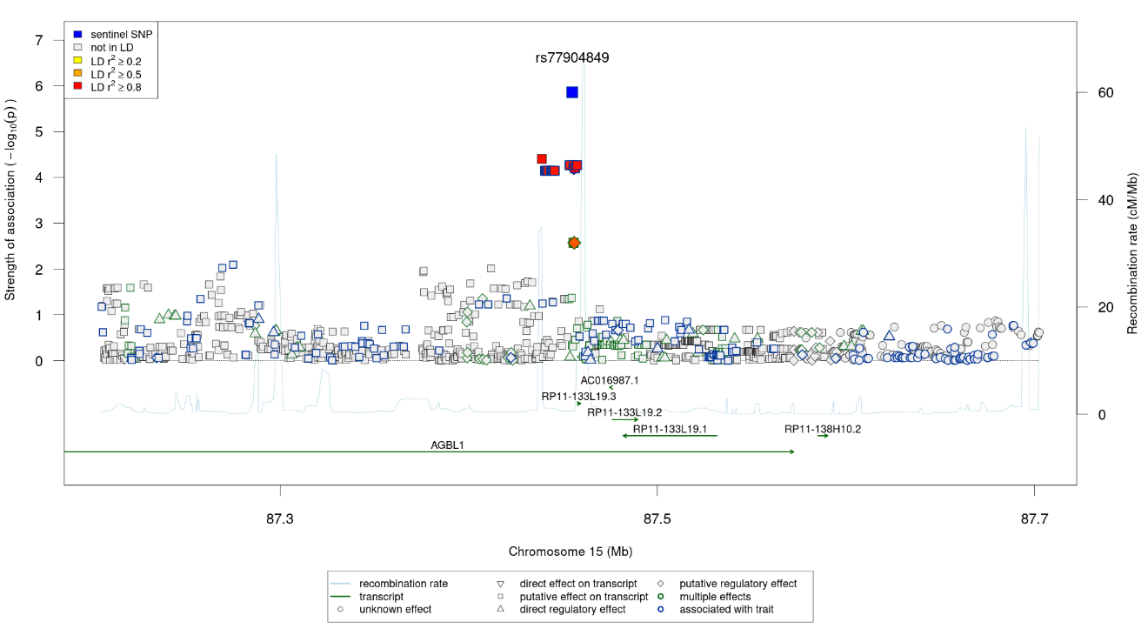

rs2034983 at chr8p23.1

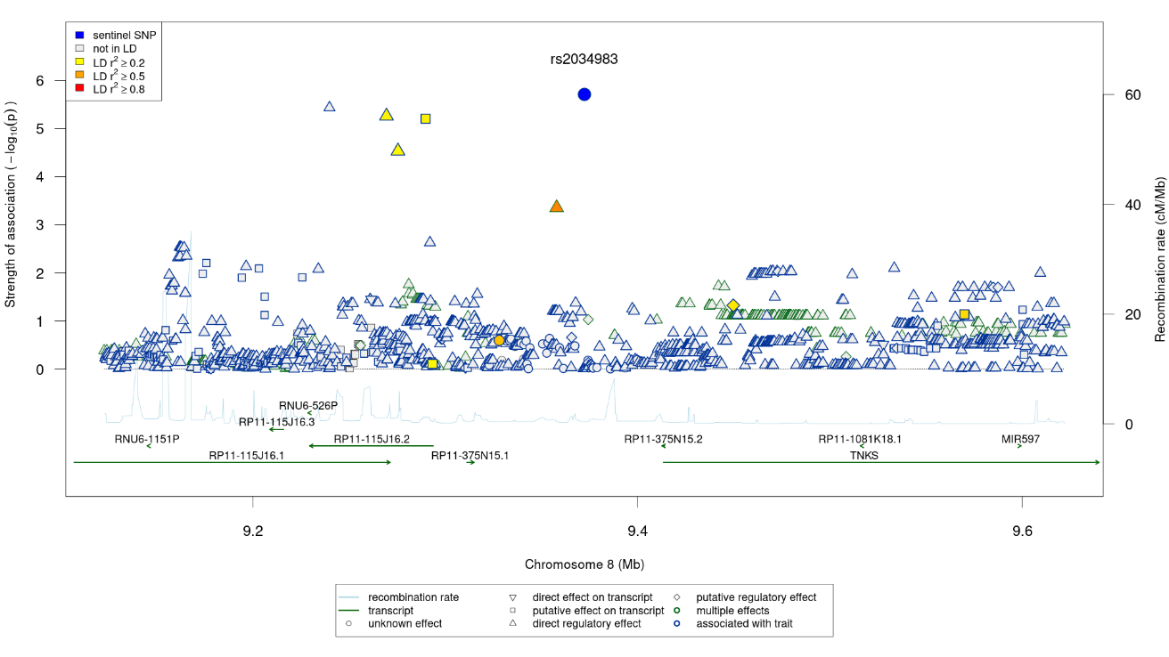

Supplementary Figure 4. Haplotype block structure (HaploView) of the *DNAJC13* locus. Block-by-block tag markers are indicated with the triangular pointers. Color coding scheme: white: D' < 1, LOD < 2; shades of red: D' < 1, LOD ≥ 2; pink: D' = 1 LOD ≥ 2.

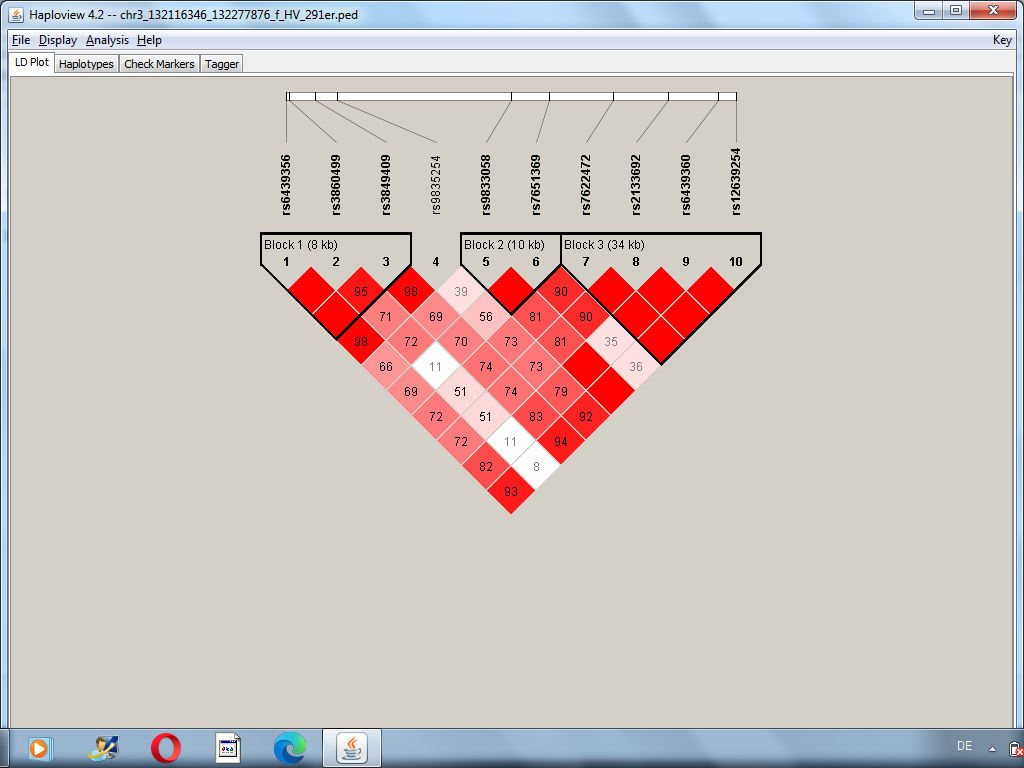

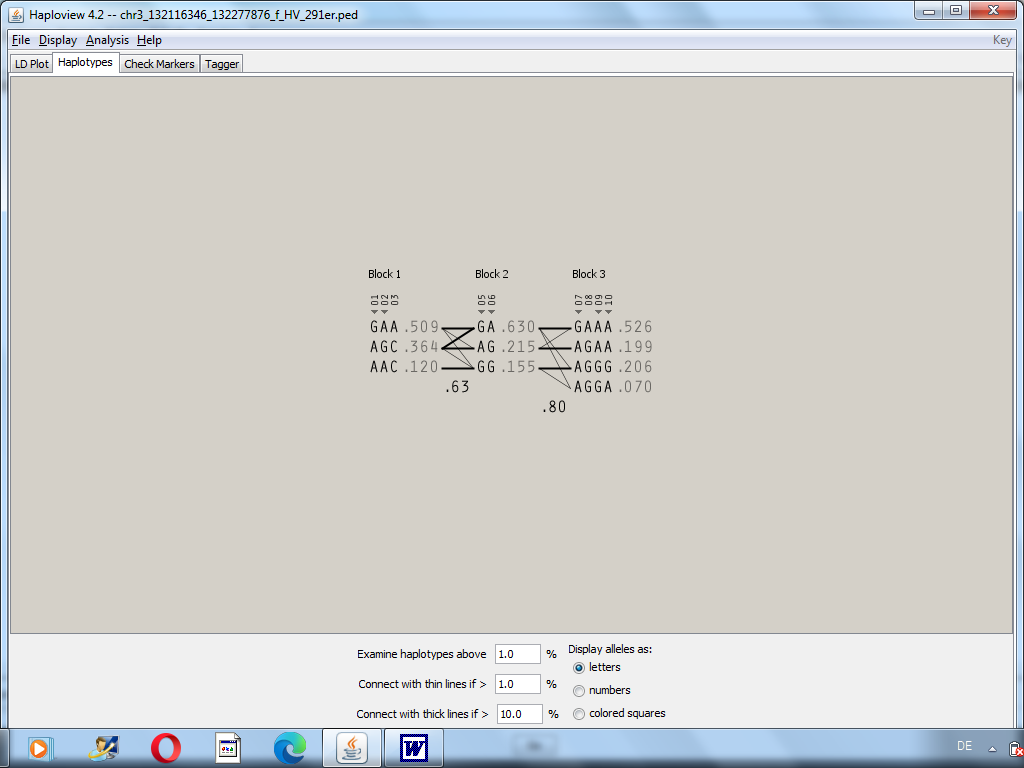

**SUPPLEMENTARY TABLES**

Supplementary Table 1. Most significant results (p < 1E-05) from single marker association analyses related to *a priori* regions of interest. a) R-VTA; b) L-VTA; c) R-NAcc; d) L-NAcc. Please see Excel-File Supplementary Table 1.xlsx.

Supplementary Table 2. Association analysis of the top ten variants from Table 1 of the main text, with principal components 1 & 2 included as covariates.

| **L-VTA** | | **R-VTA** | |
| --- | --- | --- | --- |
| **SNP** | **P** | **SNP** | **P** |
| rs57131074 | 2.5943e-06 | rs141727949 | 2.92862e-08 |
| rs10082442 | 7.34862e-07 | rs78702244 | 5.36263e-08 |
| rs7917372 | 8.53392e-07 | rs79521265 | 5.36263e-08 |
| rs1541010 | 8.53392e-07 | rs143265478 | 5.36263e-08 |
| rs1657381 | 1.06923e-06 | rs150273563 | 1.46809e-07 |
| rs1777670 | 2.54214e-06 | rs115823871 | 2.00665e-07 |
| rs141727949 | 1.3187e-06 | rs113408797 | 1.0432e-07 |
| rs150273563 | 2.6273e-06 | rs78240150 | 1.89425e-07 |
| rs78702244 | 1.64734e-06 | rs62291499 | 3.17124e-07 |
| rs79521265 | 1.64734e-06 | rs62291501 | 3.17124e-07 |
| **L-NAcc** | | **R-NAcc** | |
| **SNP** | **P** | **SNP** | **P** |
| rs12513164 | 4.33116e-07 | rs72998174 | 3.22127e-07 |
| rs78338621 | 9.80509e-07 | rs61264610 | 3.22127e-07 |
| rs34879896 | 1.58757e-06 | rs61694127 | 3.22127e-07 |
| rs9681769 | 8.83388e-07 | rs57281241 | 3.22127e-07 |
| rs12507645 | 2.62195e-06 | rs6760958 | 3.22127e-07 |
| rs12509705 | 2.62195e-06 | rs6718835 | 3.22127e-07 |
| rs77904849 | 2.0276e-06 | rs111255289 | 3.22127e-07 |
| rs75268643 | 3.66084e-06 | rs112885698 | 3.22127e-07 |
| rs2034983 | 4.99928e-06 | rs10262834 | 5.12574e-07 |
| rs12508164 | 6.38079e-06 | rs58059218 | 3.30362e-07 |

Supplementary Table 3. Haplotype association analysis (PLINKv1.07). Five tag-markers from block 1 (marker 1 [rs6439356], marker 2 [rs3860499]) and from block 3 (marker 7 [rs7622472], marker 9 [rs6439360], marker 10 [rs12639254]) were used to build the five-marker haplotypes (cf. Supporting Figure 1); F: haplotype frequency; BETA (effect size): regression coefficient; STAT: test statistic (T from Wald test); P: asymptotic p-value.

| **HAPLOTYPE** | **F** | **BETA** | **STAT** | **P** |
| --- | --- | --- | --- | --- |
| AGAGG (HAP1) | 0.0938 | 1.58 | 27.9 | 3.21E-007 |
| AAAGG (HAP2) | 0.104 | 0.108 | 0.145 | 0.703 |
| AGAGA (HAP3) | 0.0349 | 0.0548 | 0.0102 | 0.92 |
| AAAGA (HAP4) | 0.0148 | 0.0607 | 0.00602 | 0.938 |
| GAAGA (HAP5) | 0.0204 | -0.0389 | 0.0029 | 0.957 |
| AGAAA (HAP6) | 0.16 | 0.0904 | 0.118 | 0.732 |
| GAAAA (HAP7) | 0.039 | -0.196 | 0.136 | 0.712 |
| AGGAA (HAP8) | 0.0832 | 0.0685 | 0.0381 | 0.845 |
| GAGAA (HAP9) | 0.443 | -0.648 | 12 | 0.000648 |

**SUPPORTING REFERENCES**

Arnold M, Raffler J, Pfeufer A, Suhre K and Kastenmüller G. SNiPA: an interactive, genetic variant-centered annotation browser. Bioinformatics 2015;31(8):1334-1336.
